# Stem cell mechanoadaptation - Part A - Effect of microtubule stabilization and volume changing stresses on cytoskeletal remodeling

**DOI:** 10.1101/2024.07.28.605421

**Authors:** Vina D. L. Putra, Kristopher A. Kilian, Melissa L. Knothe Tate

## Abstract

Here we report on the first part of a two-part experimental series to elucidate spatiotemporal cytoskeletal remodeling, which underpins the evolution of stem cell shape and fate, and the emergence of tissue structure and function. In Part A of these studies, we first develop protocols to stabilize microtubules exogenously using paclitaxel (PAX) in a standardized model murine embryonic stem cell line (C3H/10T1/2) to maximize comparability with previous published studies. We then probe native and microtubule stabilized stem cells’ capacity to adapt to volume changing stresses effected by seeding at increasing cell densities, which emulates local compression and tissue template formation during development.

Within the concentration range 1 – 100 nM, microtubule stabilized stem cells maintain viability and reduce proliferation. PAX-stabilization of microtubules is associated with increased cell volume as well as flattening of the cell and nucleus. Compared to control cells, microtubule stabilized cells exhibit thick, bundled microtubules and highly aligned, thicker and longer F-actin fibers, corresponding to an increase in the Young’s Modulus of the cell. Both F-actin and microtubule concentration increase with increasing PAX concentration, whereby the increase in F-actin is more prominent in the basal region of the cell. The corresponding increase in microtubule is observed more globally across the apical and basal region of the cell.

Seeding at increasing target densities induces local compression on cells. This increase in local compression modulates cell volume and concomitant increases in F-actin and microtubule concentration to a greater degree than microtubule stabilization via PAX. Cells seeded at high density (HD) exhibit higher bulk modulus than corresponding cells seeded at low density (LD). These data demonstrate the capacity of stem cells to adapt to an interplay of mechanical and chemical cues, i.e. respective compression and exogenous microtubule stabilization; the resulting cytoskeletal remodeling manifests as evolution of mechanical properties relevant to development of multicellular tissue constructs.

**Significance statement:** Elucidation of mechanisms by which stem cells adapt across length and time scales may prove enabling for the development of regenerative medicine therapies and devices that emulate natural processes. Dynamic cytoskeletal remodeling underpins the emergence of structure-function relationships at the tissue length scale. Here we stabilized the tubulin cytoskeleton exogenously using paclitaxel (PAX), a microtubule depolymerization inhibitor. We probed stem cell mechanoadaptation by seeding at increasing density to introduce local compression to cells. Changes in cytoskeletal architecture and concentration of F-actin and tubulin per cell occurred in a PAX concentration-dependent manner. Compression from increasing seeding density modulated this PAX-induced cytoskeletal remodeling and mechanical properties of the multicellular constructs. Hence, mechanical cues counterbalance concentration-dependent effects of exogenous chemical microtubule stabilization.

## Introduction

Biomechanical and biophysical cues play a ubiquitous role in the emergence of structure and function in developing tissues^1,2^. Over the time and length scales relevant to processes of tissue development, stem cells use their force sensing and generating capacities to integrate and adapt to their mechanical environment, which is defined by e.g. the mechanical properties of neighboring cells and extracellular matrix (ECM)^3^. This process relies on the regulation of the cells’ cytoskeletal dynamics, i.e. F-actin and tubulin de-/polymerization^2,4,5^. The mechanisms by which cytoskeletal remodeling results in emergent functional behavior, referred to as **mechanoadaptation**^5^, is highly relevant to both tissue engineering and regenerative medicine contexts.

The use of small molecules targeting cytoskeleton de-/polymerization has proven useful for elucidating the role of the cytoskeleton in the regulation of cell shape and mechanical properties underpinning cell lineage specification^1^. Paclitaxel (PAX) inhibits tubulin **depolymerization**, thus stabilizing microtubules, increasing microtubule bundling, and leading to an increase in cell stiffness^6^. F-actin and tubulin act in concert and exhibit compensatory effects in the cell; hence, inhibition of microtubule depolymerization with PAX is accompanied by significant changes in F-actin, resulting in dose-dependent decreases in MSC migratory capacity and differentiation potential^7^. In contrast, both inhibition of F-actin **polymerization** by Cytochalasin D (CytD), as well as F-actin stabilization by jasplakinolide, reduce the concentration and organization of F-actin, resulting in a concomitant reduction of Young’s modulus in mesenchymal stem cells (MSCs), with downstream effects on MSC adaptation to mechanical stimuli and lineage commitment^8^. Use of these small molecules to specifically target tubulin and F-actin de-/polymerization, with controlled delivery of mechanical cues, provides a novel means to probe the role of the cytoskeleton in stem cell mechanoadaptation and tissue neogenesis.

F-actin and tubulin-comprising microtubules serve as the respective tension and compression resisting cytoskeletal elements of the cells’ structural scaffolding^9^. F-actin and tubulin filaments cooperatively transduce mechanical signals into bio-physical (conformational changes)^10^ and chemical signals in the nucleus^11^. Once transduced to the nucleus, these biophysical and chemical signals dictate gene regulation, and in turn, coordinate the remodeling of cytoskeletal proteins and ECM. Hence, F-actin and tubulin regulate the properties of a cell’s own tissue habitat^5,9^.

*In vivo* cells adapt to a range of volume– and shape-changing stresses, i.e. compression and shear, which embody physiological stresses intrinsic to growth, development, and maintenance of tissues^5,12^. Advances in tissue culture methods have enabled delivery of controlled volume– and shape-changing stresses, to probe how such physiological stresses might perturb a cell’s structure and function^13–16^. *In vitro*, both cell seeding density as well as mode of achieving density (seeding at versus proliferating to target density) exert significant effects on local compression, thereby influencing cell and nuclear morphometrics. Force-mediated modulation of nuclear morphology alters baseline gene expression of early mesenchymal condensation markers characteristic of skeletogenesis^13^.

In Part A of this study, we hypothesize that stabilization of microtubules via exogenous PAX exposure decreases their depolymerization and modulates the remodeling capacity of both the microtubule and the F-actin cytoskeleton, changing the associated mechanoadaptation of stem cells to changes in baseline stress state. To assess quantitatively this spatiotemporal remodeling, we measured spatial distribution of the cytoskeleton, changes in cell and nucleus volume, as well as cell shape and stiffness, over time periods relevant to early developmental processes. By quantifying the time and concentration-dependent changes in cell viability, proliferation, and cytoskeletal reorganization, we elucidated stem cells’ mechanoadaptation capacity in contexts mimicking those which occur during early stages of tissue development.

## Results

### Defining PAX concentration to maintain cell viability while challenging mechanoadaptation

#### Viability

We exposed C3H10T1/2 cells to concentrations of PAX ranging from 1 to 100 nM and measured viability using fluorometer-based measures. No significant changes in cell viability were measured with PAX exposure in the nanomolar concentration range (1 – 10 nM), neither at 24 (Fig. 1A), 48 (Fig. 1B) nor at 72 (Fig. 1C) hours after seeding. At 100 nM PAX concentration, cell viability showed no significant difference to baseline controls at 24h after seeding, but viability was significantly reduced at 48 and 72h after seeding.

**Figure 1.**
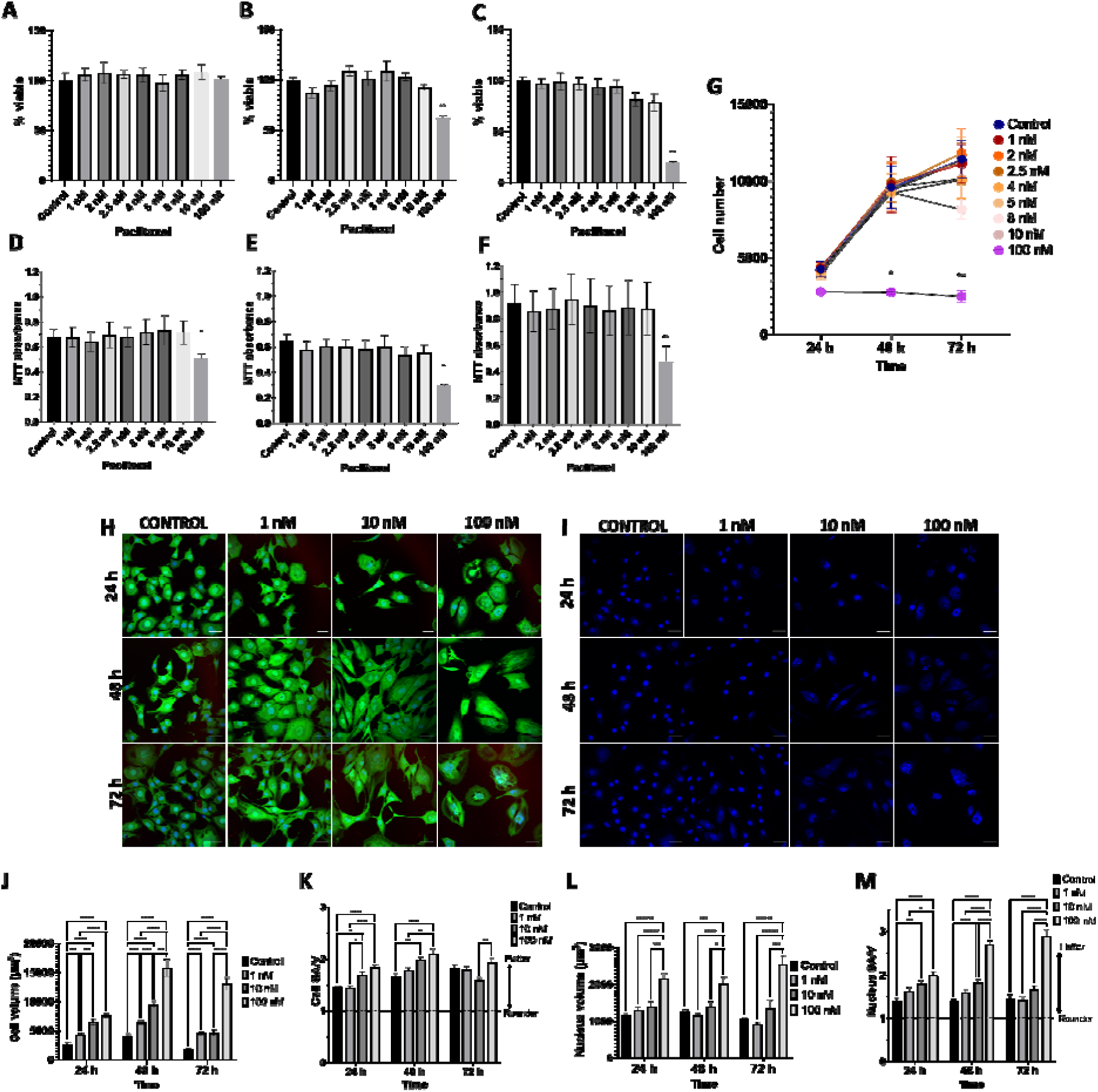
Concentration-dependent response to PAX and its effect in modulating C3H/10T1/2 murine embryonic stem cell proliferation, as well as cell and nucleus volume, shape and/or size. Cell viability measured using the Cyquant assay, after exposure to PAX ranging from 1 to 100 nM, where cell viability was maintained after 24 h (A), and significantly decreased at 48 (B), and 72 h (C) with 100 nM PAX. Metabolic activity measured via MTT assay revealed a significant reduction in MTT absorbance across the 24 (D), 48 (E) and 72 h (F) PAX exposure at 100 nM. All values are normalized to the control (DMSO treated). (G) Cell proliferation measured against the standard curve of known cell number used in the Cyquant assay. (H) Micrographs of cells (green) and their nuclei (blue) demonstrate gross and increasing changes in cell and nucleus size and shape, as well as nuclear fragmentation, with increasing PAX concentration and increasing cell number with increasing time in culture. Further, for each PAX concentration studied, cell volume increased from 24 to 72 h in culture after which cell volume decreased relative to 48h but remained significantly higher than cell volume at 24 h (scale bar = 50 μm). (I) PAX-induced changes in nucleus morphology include formation of multinuclear fragments and flatter morphology. (J) Quantification of cell volume (V) and surface area (SA) and shape (SA/V) demonstrates effects of increasing PAX concentration and time in culture. At 24, 48, and 72h a linear increase in cell volume was observed with each tenfold increase in PAX dose (Y = 31.00*X + 4687, Y = 92.97*X + 6683, Y = 96.75*X + 3405). (K) Cell shape is defined as SA/V ratio, where relatively rounder cells have SA/V close to 1 and flatter cells with SA/V above 1. At 24 and 48h, cells exhibit concentration dependent flattening which stabilized at 72h. (L) Nucleus volume increases also occur in a time and concentration dependent manner, concomitant to cell volume increase. (M) Nucleus SA/V is visibly higher with increasing PAX concentration over time indicative of their flatter and wider shape. Error bars represent ± standard error of mean. Significant differences are presented between groups (**** p < 0.0001, ** p < 0.01, * p < 0.05) and analyzed with one-way or two-way ANOVA repeated measures with Tukey’s multiple correlation test).

In addition, we used the MTT assay of metabolic activity as an indirect measure of PAX cytotoxicity. Metabolic activity showed no significant change with PAX exposure at any timepoint measured except for the cells treated with 100 nM PAX, which exhibited a significant decrease in metabolic activity at all timepoints measured (24 (Fig. 1D), 48 (Fig. 1E) and 72 (Fig. 1F) hours after seeding).

Hence, neither cell viability nor metabolic activity were affected significantly by exposure to PAX concentrations up to 100 nM, where a tipping point was observed. Namely, exposure to 100 nM PAX was associated with decreasing metabolic activity, and decreasing viability after 48 and 72h of culture; hence, **we chose 100 nM PAX as an upper bound for probing cell adaptation and survival to exogenous microtubule stabilization via PAX.**

#### Proliferation

Changes in cell proliferation were measured using a Cyquant fluorometer, by extrapolating cell number from standard curves of known cell numbers, for all microtubule stabilized groups at 24, 48 and 72 h after seeding (Fig. 1G). Below 100 nM PAX, the observed proliferation rate was similar to that of the control group. A significant fall in cell number was evident at 72 h in both the 8 nM and 10 nM PAX groups. At the upper bound, 100 nM PAX concentration, cell number remained low and continued to decrease up to 72 h.

#### Live/dead imaging and morphological changes

Morphological metrics were acquired during live/dead imaging of microtubule stabilized stem cells’ time– and concentration-dependent decreases in cell viability and proliferation.

Over 72 h of microtubule stabilization, a majority of cells exhibited the calcein AM label, indicative of maintained viability (Fig. 1H). However, the number of viable cells per field of view decreased with time and with increasing PAX concentration, which contrasted with observations of the control group that reached greater than 80% confluency at 72 h (Fig. S1A). The cells exposed to 1 nM PAX demonstrated similar proliferation rates as control cells; at higher concentrations, e.g. 10 nM and 100 nM PAX, the cell number decreased over time.

In general, with increasing PAX concentration and exposure time, cells exhibited increased volume with flatter cell and nucleus shape (measured by SA/V, surface area to volume ratio). Nuclear changes occurred concomitant to changes observed in cell structure, including a concentration-dependent increase in nuclear volume and fragmentation (Fig. 1I) at the highest PAX concentration tested (100 nM). In contrast the control and lower concentration (below 100 mM) microtubule stabilized cells exhibited round and intact nuclei. Microtubule stabilized cells grew significantly in volume, in a time– and concentration-dependent manner, with the upper bound, 100 nM group showing a significantly higher volume increase than all other groups at 48 and 72 h (Fig. 1J, Fig. S1B).

Cell shape measures, defined by the surface area to volume ratio (SA/V), indicate relative flattening of cells and/or nuclei with higher relative SA/V and relative rounding of cells and respective nuclei, with lower SA/V. To measure changes in shape independent of cell volume, the cell and nucleus SA/V were normalized to the SA/V of a perfect sphere (3/R) with the same volume, according to previous protocols^13^. The normalized SA/V with value of 1 indicates perfect sphericity (roundness) and above 1 indicates a flatter and more spread shape. At 24 and 48 h exposure to the microtubule stabilizer PAX, cells exhibited concentration-dependent increases in SA/V, i.e. flattening, while at 72h all microtubule stabilized groups showed similar and higher SA/V ratios (Fig.1K). The nucleus volume increases paralleled the PAX concentration-dependent cell volume increases (Fig. 1J and Fig. S1C). Compared to changes in cell shape, the flattening of nuclei (relative increase in SA/V) was more prominent and increased with microtubule stabilization time and PAX concentration (Fig. 1K). The increase in cell/nuclei SA/V due to microtubule stabilization is modelled in Fig. S1D by changing the one of the axes lengths (either x, y, or z) of a sphere to show the elongated and flattened cell/nuclei shape.

Over time, cell thickness (measured in the z-direction or distance from basal to apical, Fig. S1E) decreased in culture, except for the 100 nM PAX concentration group where cell thickness remained constant over time (Fig. S1F), accounting for the effect of exogenous microtubule stabilization on the increase in nucleus size. Nuclear thickness measurements (Fig. S1G) were consistent across the PAX concentration groups but reduced over time in parallel with reduction of cell thickness.

### Effect of exogenous microtubule stabilization on cell and subcellular remodeling

Once we identified the PAX concentrations that maintain viability while challenging cell mechanoadaptation, we probed cytoskeletal remodeling and cell mechanoadaptation in response to exogenous microtubule stabilization. In general, with inhibition of microtubule depolymerization, cells increased in size concomitant to cytoskeletal remodeling which was grossly evident and paralleled the time– and PAX concentration-dependent changes in cell shape and increases in cell volume, as described in further detail below (Fig. 2A).

**Figure 2.**
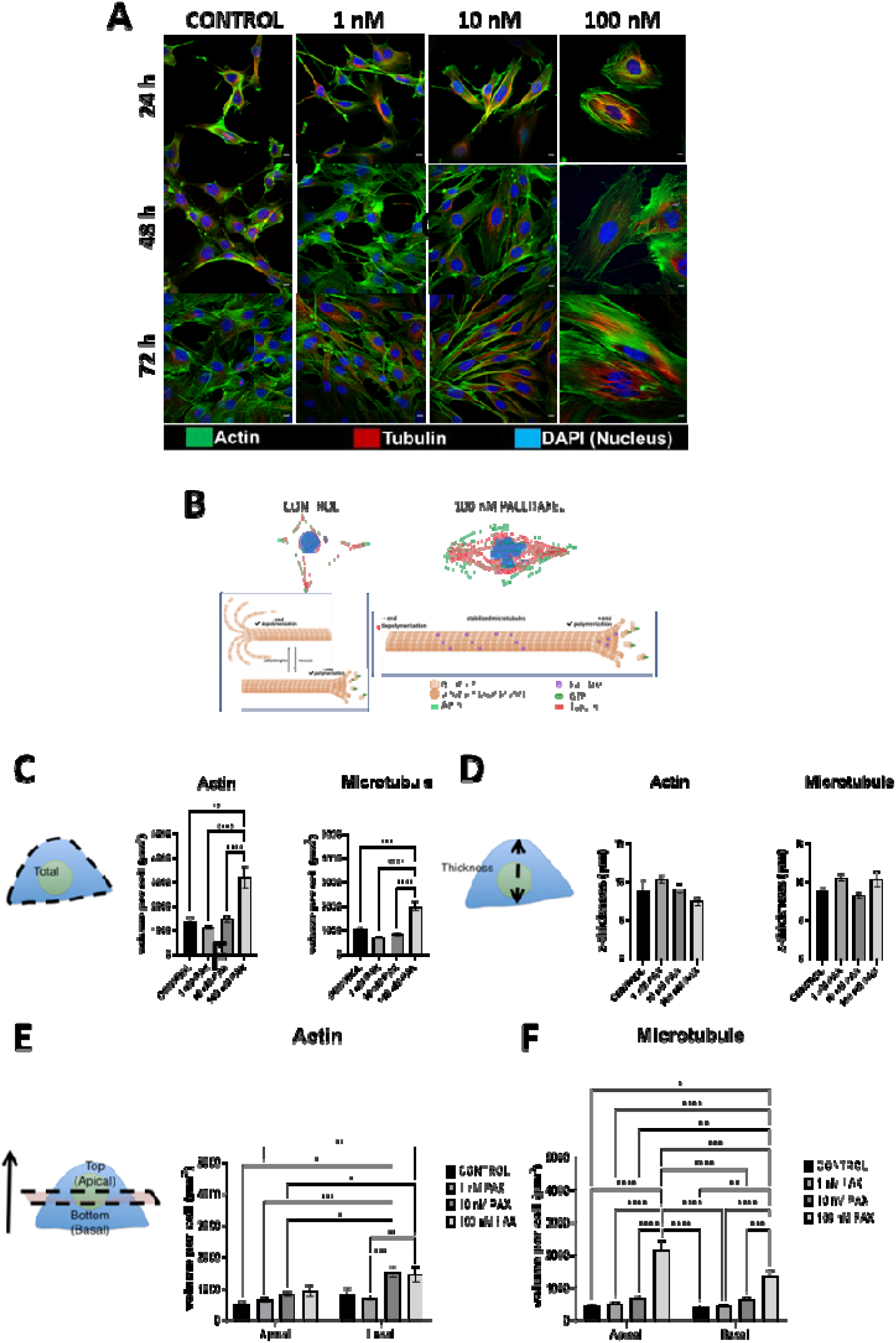
With increasing concentration of microtubule depolymerization inhibitor, PAX, microtubule and actin dynamics and spatial organization exhibit anisotropic and disparate effects. (A) Confocal images of actin (green), microtubules (red) and nuclei (blue) demonstrate structural changes in the respective cytoskeletal elements’ length and organization (angle of orientation in space including alignment in the focal plane and distribution in space, as well as higher order architecture). (B) Schematic of cytoskeleton remodeling due to PAX. In unexposed cells, microtubule polymerization and depolymerization occur at an equal rate, thus maintaining the balance of pushing and pulling forces necessary to carry out protein transport and cell division. In PAX exposed cells, microtubules are stabilized and depolymerization is inhibited, preventing further cell division. The continuation of microtubule polymerization results in a pushing force towards the cell periphery and increased cell volume. (C) Total actin and microtubule volume per cell are significantly higher in the 100 nM PAX exposed cells which show three– and two-fold respective increases. No significant differences in total actin and/or microtubule were observed between control cells and microtubule stabilized cells, i.e. exposed to less than 100 nM PAX. (D) Thickness of the actin and microtubule were not significantly different between the control and microtubule stabilized groups. (E) Upon PAX exposure the spatial distribution of actin was significantly higher in the basal region but not in the apical region. (F) In contrast, both the apical and basal distribution of microtubules were significantly higher in 100nM PAX group than in all other groups. Scale bar: 10 μm. Error bars represent ± standard error of mean. Significant differences are presented between neighboring values (**** p < 0.0001, *** p < 0.001, ** p < 0.01, * p < 0.05) from non-parametric, two-way ANOVA and Tukey’s multiple comparison.

#### Cytoskeletal remodeling – amount and spatial distribution, and orientation of F-actin and tubulin

Based on observation of confocal images (Fig. 2A), control cells showed short peripheral F-actin arcs, typically emanating in multiple directions, and balancing protrusion and adhesion to neighboring cells and/or the substrate. In contrast, PAX-exposed cells lacked such F-actin arcs; instead exhibiting thicker, longer and more pronounced F-actin stress fibers than control cells; these stress fibers were oriented along the highly concentrated unidirectional growth of the depolymerization inhibited microtubules (Fig. 2B). In addition, thicker bundles (relative to controls) of microtubules aggregated at high density in the perinuclear region and between nuclear fragments (Fig. 2A,B).

Quantitative measures showed significant increases in the total concentration of F-actin and microtubules per cell in the 100 nM PAX exposed group compared to the control and other lower concentration PAX groups (1, 10 nM), corroborating gross observations of PAX concentration-dependent F-actin alignment (Fig. 2C; Table S2). Despite this, the cytoskeleton thickness spanning the cell height remained similar across all microtubule stabilized and control groups (Fig. 2D).

To quantify the spatial distribution (emergent anisotropy) of the F-actin and microtubule as a function of PAX-concentration, cells were segmented into apical and basal regions (Fig. S1E). For the 10 nM PAX exposed group, significantly more F-actin was observed in the basal region of the cell. For the 100 nM PAX exposed group, a significant increase in microtubule was observed in the apical region of cells and a (not significant) increase in F-actin was observed in the basal region of the cell. Together, these observations confirmed emergent anisotropy in spatial distribution of both F-actin and microtubule filaments with exposure to 10nM and 100 nM PAX (Fig. 2E). While the increase in F-actin concentration was more prominent in the basal region of the cell for the 10 and 100 nM PAX – microtubule stabilized groups, the respective increase in microtubule concentration was observed in the apical region of the 100 nM PAX exposed cell, indicative of reciprocity or cross talk (Fig. 2F).

Our working hypothesis was that the coexistence and orientation (see gross observations above and quantification below) of F-actin and microtubules remodeling indicated cytoskeletal cross talk conducive to balancing of forces at cell boundaries (see Discussion). To test this hypothesis mechanistically a cohort of cells was treated with CytD, which inhibits F-actin polymerization. In contrast to effects observed with PAX exposure, which inhibits microtubule depolymerization, inhibition of F-actin polymerization with CytD drastically reduced cell spreading, viability (Fig. S2A-C), proliferation (Fig. S2D), metabolic activity (Fig. S2E-G), and cell volume (Fig. S2H-J). Inhibition of F-actin polymerization also removed tension that F-actin normally generates in adherent cells, leading to collapse of stress fibers and shortening of F-actin branches with increasing CytD concentration (Fig. S2K). The immediate (within a few hours) but reversible effect of CytD on F-actin stress removal and build-up, made it challenging to observe stem cell adaptation at later time points when multicellular constructs emerge at the tissue length scale.

### Microtubule stabilization modulates cytoskeletal remodeling and influences cell mechanical properties in both adherent and suspended cells

#### Cytoskeletal alignment

The PAX concentration-dependent degree of alignment of the cytoskeletal network was quantified using Noise Based Segmentation (NoBS, see Methods, Fig. 3A). NoBS visualization and metrics for F-actin showed increasing alignment in the global stress fibers detected in the overall field of view and as a function of increasing PAX-concentration. Simultaneous measurement of F-actin alignment with increasing NoBS working radius (from 1 to 100 μm) within a field of view revealed an optimum alignment at a radius of 15 μm, corresponding to the average radius of adherent C3H/10T1/2 cells (Fig. 3B). With 100 nM PAX exposure, a majority of the detected F-actin filaments were continuous even beyond the 15 μm distance, which contributed to the measured high alignment score. In contrast, the low alignment score measured in control cells resulted from discontinuous fibers detected within the same radial distance. When 3D images for each group were aligned using the mid-Z plane of the nucleus as a reference for each image (marked at 0 μm dotted line), a unique F-actin alignment profile was observed in the basal and apical region of microtubule stabilized cells (Fig. 3C). Furthermore, there was a significant increase in F-actin alignment in the basal compared to the apical regions of the 100 PAX-exposed as well as control cells. This is remarkable when one considers that the increase in F-actin concentration was higher in the apical region of the cell for this group, suggesting remodeling processes occur in the basal region prior to the apical region (see Discussion).

**Figure 3.**
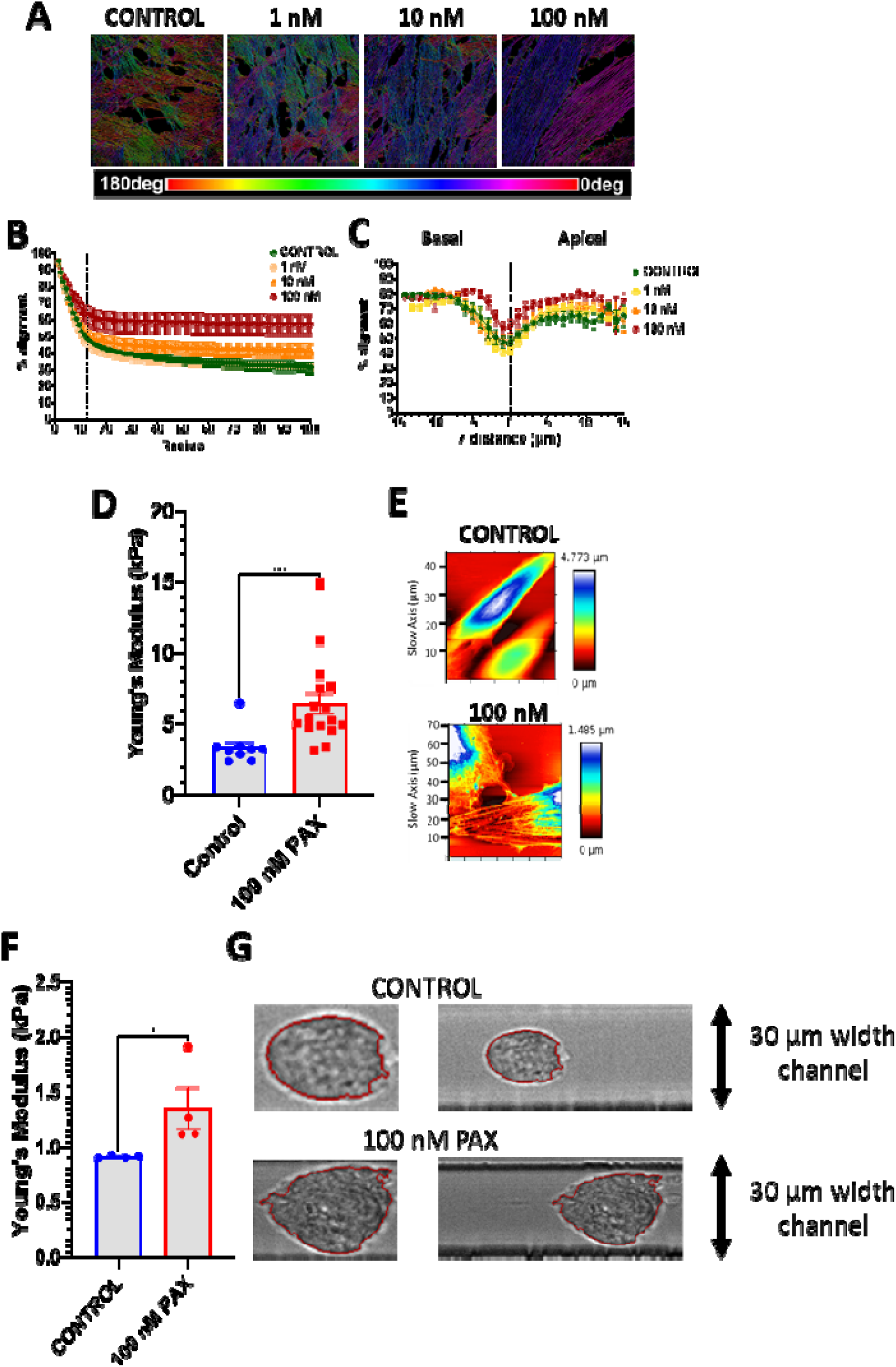
F-actin alignment increases with increasing PAX concentration and a significant increase in cell stiffness. Actin orients in a predominant direction (0 – 180 degrees), as measured by Noise based Segmentation (NoBS) and depicted on a color scale (A) and quantitatively at laser scanning radii of 1 – 100 μm (B). Alignment increases with increasing PAX concentration, with significant effects above 10 nM PAX concentration. (C) Actin alignment varies from the basal to the apical surface of the cell, with higher alignment in the basal region (up to 10 μm below the mid-plane) than in apical region (up to 20 μm above the mid-plane) with microtubule stabilization. Nucleus z-thickness is used to determine the mid-plane of the cell (dotted line marked at 0 μm) and to define the apical and basal regions. (D) AFM analysis of the mean Young’s modulus of microtubule stabilized cells, where exposure to 100 nM PAX for 72 h resulted in twice stiffer cells compared to the control cells. Each point depicts the mean stiffness of a single cell. (E) The height map of the cells (indicating the lower and higher topographies), determined from the contact point of each indentation measurement, reveals that stress fibers of microtubule stabilized cells appear thicker and more defined, thus contributing to the overall increase in cell stiffness. (F) Deformability cytometry of microtubule stabilized cells. After trypsinization, cells in suspension flowed through a 30 μm wide channel while measuring cell deformability and stiffness. Microtubule stabilized cells exhibit a higher Young’s modulus with (G) larger area and volume (Fig. S2A-B) compared to smaller and rounder control cells. Error bars represent ± standard error of mean. Significant differences are presented between neighboring values (**** p < 0.0001, *** p < 0.001, ** p < 0.01, * p < 0.05) from non-parametric, two-way ANOVA and Tukey’s multiple comparison.

#### Mechanical properties at the cell and subcellular length scales

To examine changes in bulk and subcellular mechanical properties of cells, cell stiffness (Young’s Modulus) was measured using both Atomic Force Microscopy (AFM) of adherent cells as well as deformability cytometry of nonadherent (trypsinized) cells.

#### AFM based stiffness measures

Control cells exhibited a mean Young’s Modulus of 3.3 kPa. PAX-exposed cells were significantly and nearly twice stiffer with a mean Young’s Modulus of 6.4 kPa (Fig. 3D). Indentation maps showed vertical deflection of the AFM tip as it probed different locations in single cells and provided spatial distribution measure of subcellular stiffness (Fig. 3E).

To examine whether the PAX-induced stiffening was independent of adhesion, cell stiffness was measured with cells in a suspended state, where cells were trypsinized and flushed through a 30 μm wide microfluidic channel for deformability cytometry. Cells in suspension were softer than respective adherent cells in control and experimental groups. Microtubule stabilized (PAX) and suspended cells were significantly stiffer than the control cells with a 2-fold increase in Young’s Modulus (Fig. 3F). Consistently, trypsinization of PAX-exposed cells was associated with a significant increase in cell area and volume (Fig. S3A,B). Microtubule stabilized cells appeared larger and with irregular contours compared to the control cells which were smaller and rounder (Fig. 3G). This contributed to the larger deformation observed with deformability cytometry (Fig. S3C), as larger cells were deformed more than smaller cells as they flowed through the channel.

Since both cell and cytoskeletal organization, as well as mechanical properties, changed with adhesion state and as a function of PAX concentration, we then probed how changing the local mechanical environment could further modulate the observed concentration– and boundary condition-(adherent vs. non-adherent) dependent changes; our approach was to control changes to the local mechanical environment of cells by increasing seeding density^4,13,17^. In a follow-on set of experiments (see [Part B manuscript]), we probed the effects of tuning substrate stiffness,^18^ with and without cell seeding density changes, on cell mechanoadaptation.

### Increasing seeding density modulates PAX-induced increases in cell volume and shape changes

Given that microtubules serve as compression resisting filaments of the cells, we next investigated how increasing seeding density, which was previously shown to induce local compression,^13,17^ modulates PAX concentration– and time-dependent changes in cell morphology and cytoskeletal concentration/spatial distribution. We seeded the cells at low (LD 5,000 cells/cm^2^), high (HD 15,000 cells/cm^2^) and very high seeding density (VHD 45,000 cells/cm^2^) and exposed to exogenous microtubule stabilization (PAX) for 72 h. Previous studies have shown that higher initial seeding density profoundly influences the respective volume and shape of cells and their nuclei, impacting the way cells experience their local mechanical environment^4,13,17^. When seeded at LD, cells can proliferate to higher density over time until confluence is reached; seeding at VHD creates a local compressive environment for cells at the time of seeding, changes cell volume differentially to nucleus shape (like a mechanical clutch), and influences cell behaviors such as lineage commitment^4,13,17^.

Using this approach to control the local environment, we quantified cell and cytoskeletal adaptation over time. Our hypothesis was that, since microtubule stabilization via PAX increases the compression resisting capacity of cells (microtubules), less volume expansion would be necessary to counteract the local compression induced by increased seeding density, *I.e.* **for a given local pressure induced by increasing seeding density, a stiffer cell (higher *E*) will better resist the local pressure without needing to balance forces at the cell boundary by increasing volume, increasing microtubule concentration, or remodeling microtubules**^5^.

#### Validation of local compression induced by seeding at increased densities

We first validated previous studies with the same model system from our group^4,13,17,19^, showing that cells experience increasing local compression with increasing seeding density, We observed multidimensionality of cells seeded at increasing density, i.e. for each seeding density studied, cells overlap each other. Due to the stochastic, spindle-like shape of cells with protrusions when adherent, intercellular space can be observed even in the most basal plane and at very high seeding density. How cells regulate this intercellular space is a property of their structure and function as multicellular systems and is intrinsic to the local compression cells sense. At the sub-cellular level, our data shows that increasing seeding density indeed modulates the increase in PAX-mediated cell volume, actin and microtubule concentration as detailed below. An enlarged 3D view of cells seeded at higher density demonstrates the negligible intercellular space (Fig. 4).

**Figure 4.**
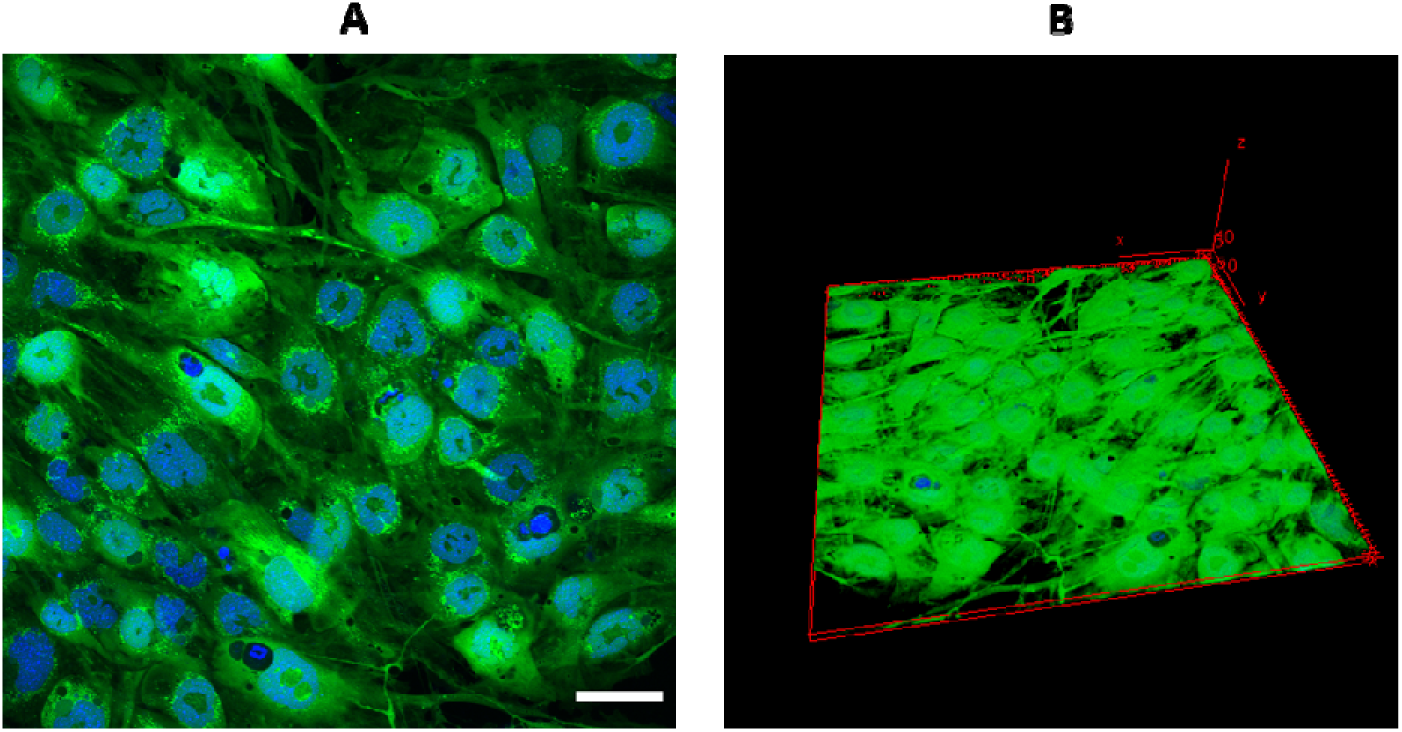
Multidimensional, multicellular constructs achieved via seeding at target seeding density. A) C3H/10T1/2 cells treated with 100 nM PAX and seeded at 45,000 cells/cm^2^, very high density (VHD). B) 3D view of the image showing multidimensionality as cells overlap each other and adapt to the local compression. Scale bar = 50 μm.

#### Impacts of local compression on cell morphology

Adaptation of cell morphology over time was evident for all control and experimental groups, with control cells increasing in volume by 1.5-fold when seeded at HD and VHD, respectively, compared to LD for 72h (Fig. 5A). An opposite effect was observed in microtubule stabilized (PAX exposed) cells, where cell volume decreased by circa 40% and 25% when seeded at HD and VHD, respectively compared to LD for 72h, with a less prominent effect at the earlier timepoints, 24h (Fig. 5B) and 48h (Fig. 5C). In general, PAX-concentration dependent cell volume increases were greatest for cells seeded at LD (Fig. 5D, Fig. S4A, B at 24 and 48 h) whereby cell volume was 5-fold higher in PAX-exposed than in control cells, compared to those at HD and VHD where PAX-mediated volume increase was only 3 to 4 –fold respectively. 3D view of cells at VHD shows overlapping of cells with maintenance of intercellular space (Fig. S5). A *post hoc* linear regression analysis of cell volume increase with increasing PAX concentration over 72h, demonstrates that cell volume increases with increasing PAX concentration, and with increasing significance at later time points (Fig. 6). Also of note, PAX-exposed cells that were trypsinized and suspended showed a more than 100% increase in volume (Fig. S3B), which contrasts with the volume decreasing effect on PAX-treated cells in an environment of increasing local compression (via seeding at increasing densities).

**Figure 5.**
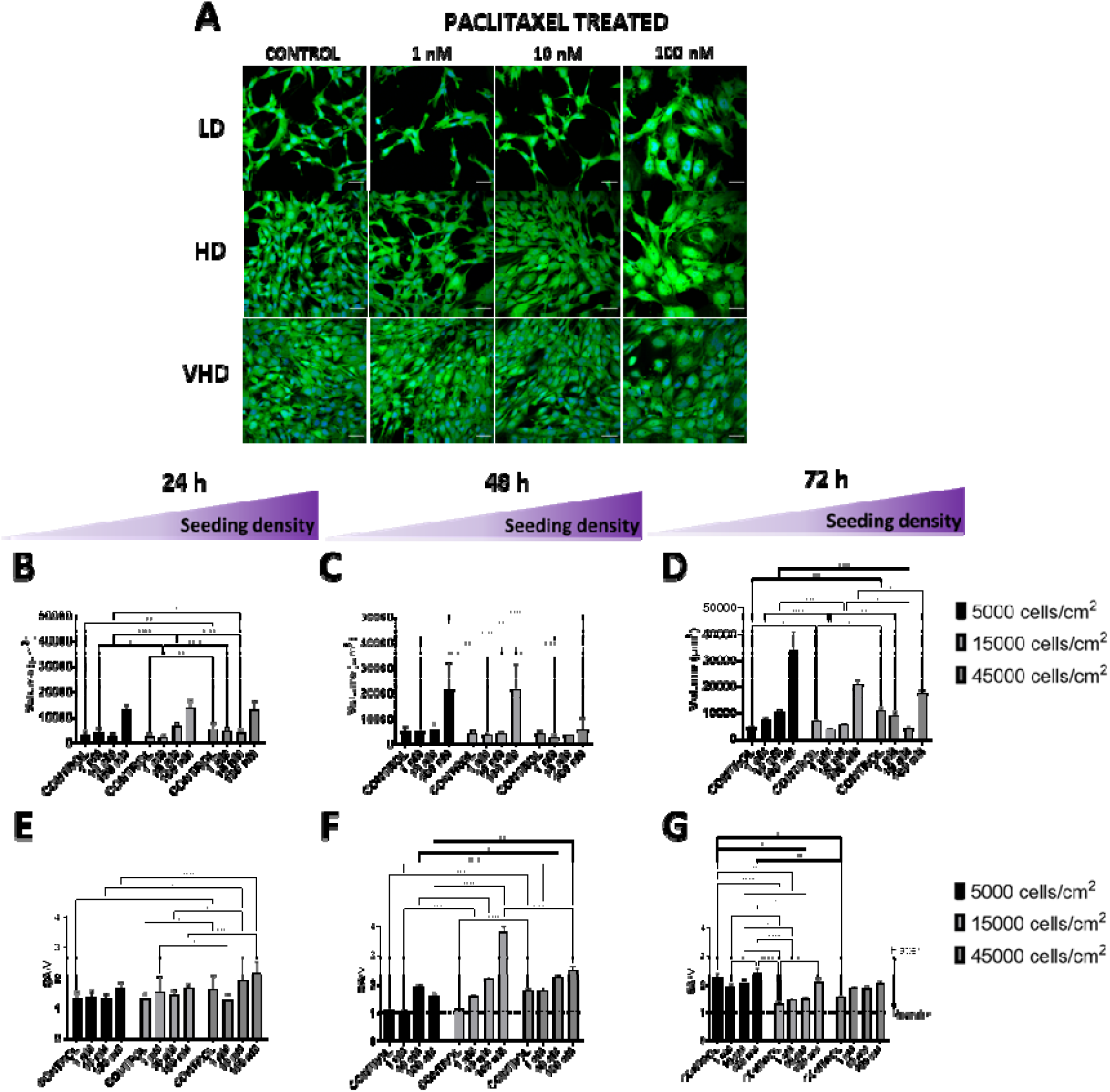
Seeding at increased density exerts a greater effect in modulating cell volume increase mediated by microtubule stabilization. Increasing seeding density, shown previously to introduce local compression^13,17^, modulates stem cell and nucleus shape/volume, as well as mechanical properties. (A) Microtubule stabilization, after seeding at low (LD), high (HD), and very high density (VHD) and culture of cells over 72h, exerts a profound influence on cell shape and volume (Scale bar = 50 μm). PAX concentration dependent cell volume increases are grossly observable after 24 h culture (B) in LD, HD, and VHD, and becoming less evident at later timepoints, i.e. 48h (C) and 72h (D). With time in culture, cells exhibit small increases in volume and cells seeded at high-density generally flatten (e.g. exhibit a higher SA/V) compared to cells seeded at low-density cells in the earlier time points 24h (E) and 48 h (F), eventually reaching a plateau at 72h (G).

**Figure 6.**
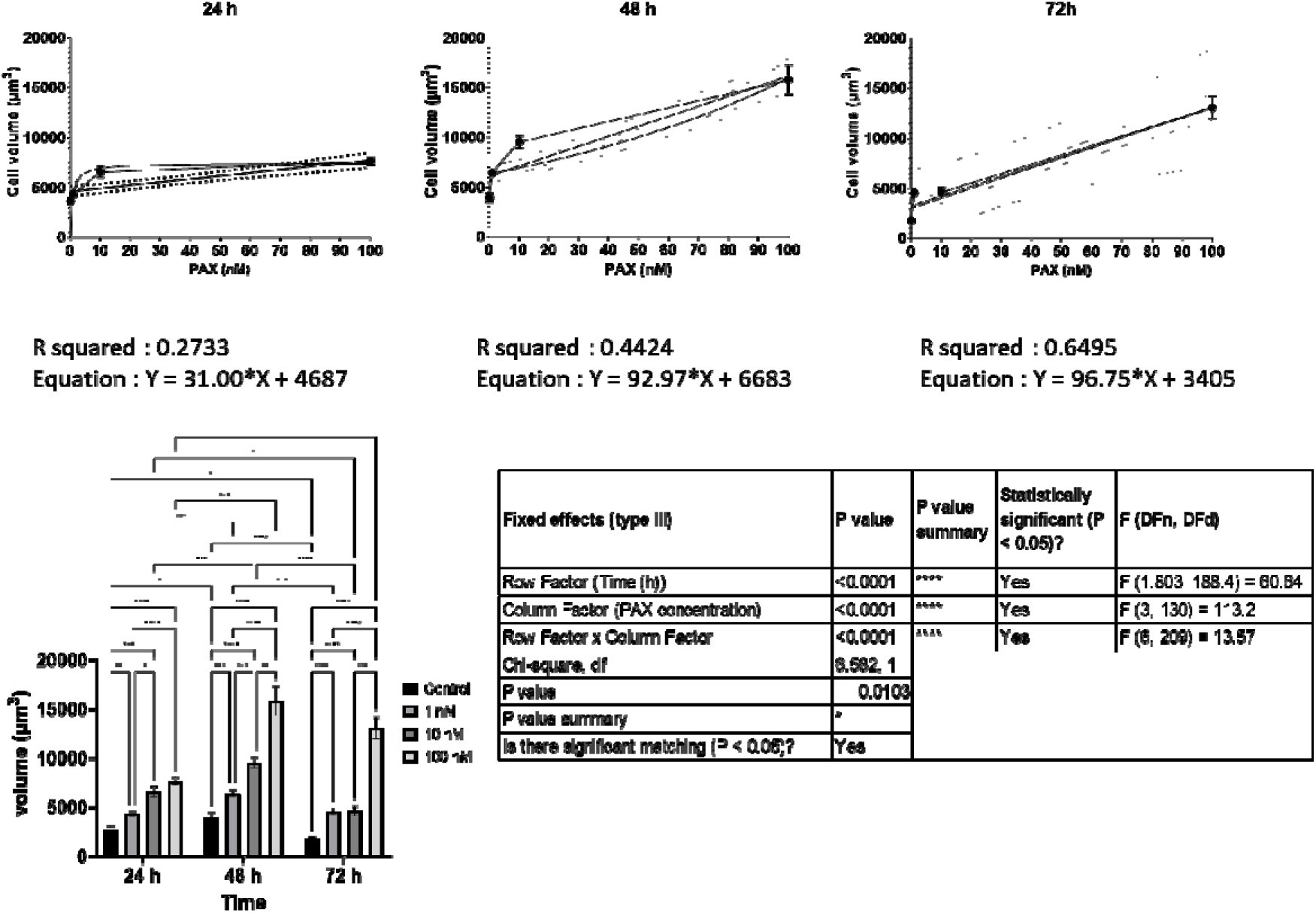
Exogenous microtubule stabilization results in increased cell volume. Top: Linear regression analysis of cell volume increase with increasing PAX concentration over 72h – showing that cell volume increases with increasing PAX concentration, and with increasing significance at later time points. Bands represent 95% confidence intervals. Error bars represent standard error of men (SEM). Bottom: Two-way ANOVA, fixed effect type III analysis of cell volume increase with increasing PAX concentration (based on Figure 2D in manuscript), with Tukey’s multiple comparison test, demonstrates that PAX-mediated cell volume increases correlate with time.

Shape changes were also evident in association with local compression, where the SA/V was slightly higher at HD, indicative of cell flattening over time as the local compression limits the cells’ volumetric growth. This contrasts with cells seeded at LD, especially microtubule stabilized cells which grow in surface area and volume at a faster rate than control cells and thus show a consistent SA/V across 72h culture (Fig. 5E-G).

#### Impact of local compression induced seeding density on nuclei

Nucleus volume showed an increase with PAX-concentration and seeding density (Fig. S4C) but not time. Increasing density at seeding moderately compensates for the PAX-induced nuclear volume increase (Fig. S4D-F). In contrast, nucleus shape (SA/V) appears to depend less on cell density where the PAX concentration-dependent cell flattening (increase in SA/V) remains similar across all time points (Fig. S4G-I).

### Increasing seeding density modulates PAX-induced increases in F-actin and microtubule concentration and their emergent anisotropy

#### Impact of seeding density on cytoskeletal concentration and spatial distribution

The influence of seeding density on the spatial distribution of F-actin and microtubule during PAX exposure demonstrates the anisotropy of both cytoskeletal filaments (Fig. 7A). Increasing seeding density does not influence F-actin or microtubule concentration across the cell z-thickness (Fig. 7B, C). Both F-actin and microtubule exhibit PAX dose-dependent increases within the total cell at LD, which is higher in F-actin (Fig. 7D) than in microtubule (Fig. 7E), which is observable to a lesser degree in cells seeded at HD. Consistently, when quantified in the apical (Fig. 7F,G) and basal region (Fig. 7H,I), the dose-dependent increase of F-actin and microtubule concentration is more notable at LD and diminishes at HD and VHD.

**Figure 7.**
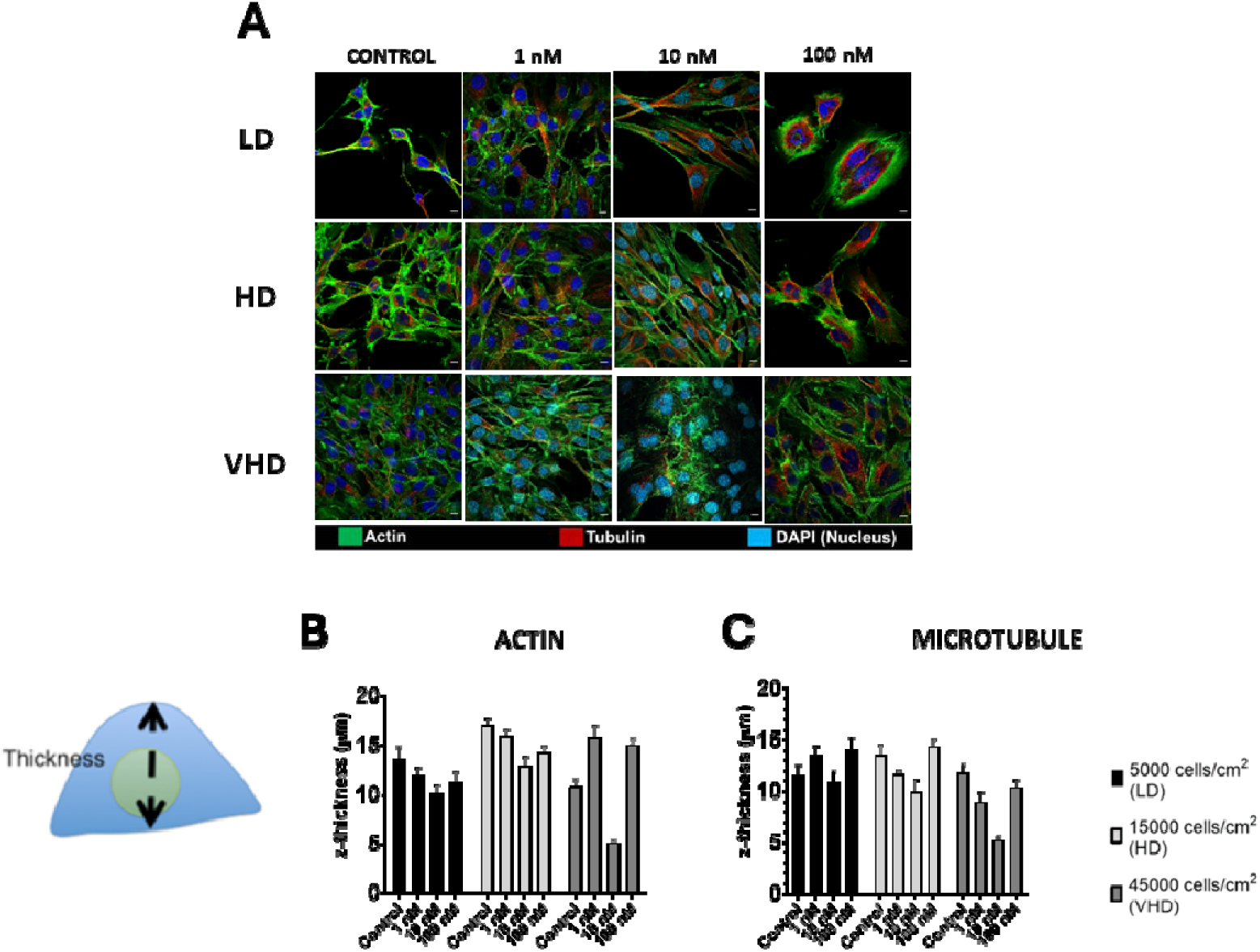
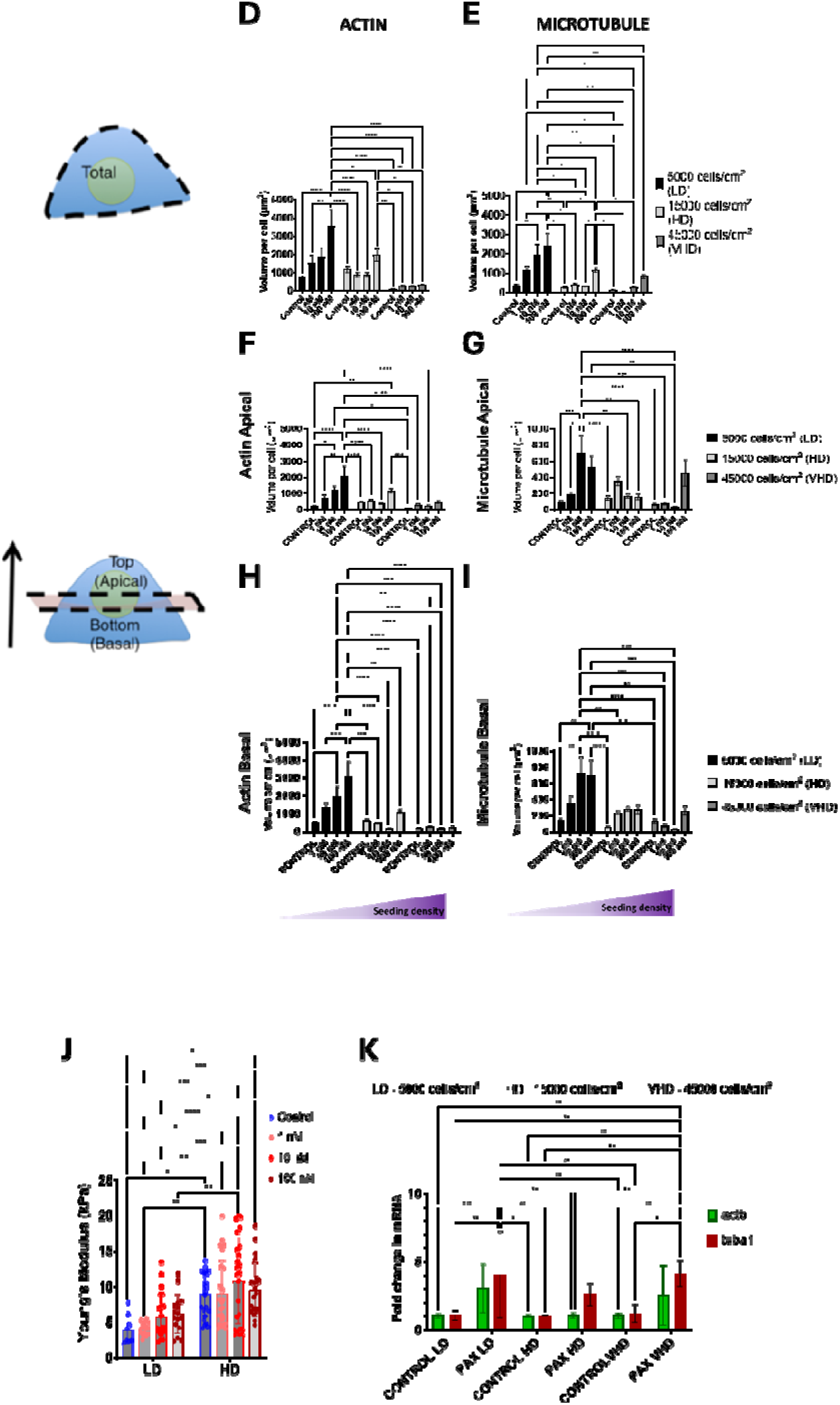
Seeding at increased density amplifies actin and microtubule emergent anisotropy and response to microtubule stabilization (PAX). (A) Initial seeding density is associated with significant effects on the concentration and spatial distribution of actin and microtubule (scale bar 10 μm), albeit no significant differences in total actin (B) and microtubule (C) thickness. The increase in total actin (D) and microtubule (E) concentration per cell is lower with increasing seeding density, so is the increase in the apical (F,G) and basal (H,I) region. PAX-concentration dependent stiffening is significant in cells seeded at low density but not in those seeded at high density or in control cells (J). (K) RT-PCR of actin and microtubule in cells seeded at low (LD), high (HD) and very high density (VHD). Error bars represent ± standard error of mean. Asterisk(s) represent significant difference at **** p < 0.0001, *** p < 0.001, ** p < 0.01, * p < 0.05, two-way ANOVA, with Tukey’s multiple comparison test.

#### Impact of seeding density on cell stiffness

Cells seeded at HD exhibited a significantly higher mean Young’s Modulus compared to cells seeded at LD. Exposure to PAX was also associated with significantly higher respective Young’s Moduli for both cells seeded at LD and HD except for cells exposed 100 nM PAX. which showed similar relative stiffnesses to 10 nM PAX exposed cells (no significant differences) and higher albeit no significant differences attributable to 100 nM PAX exposure versus control cells seeded at HD (Fig. 7J). In general, cells seeded at HD built multicellular constructs and exhibited higher bulk moduli, however, the relative increase in HD cell stiffness due to PAX exposure was smaller than in LD cells exposed to PAX.

At an mRNA level, the expression of microtubule and F-actin were consistently higher with microtubule stabilization via PAX (Fig. 7K). With microtubule stabilization, microtubule and F-actin were increased by 3-fold and 4-fold respectively in cells seeded at LD. At high (HD) and very high density (VHD), the increase in both F-actin and microtubule measured at an mRNA level were consistent with the modulation of F-actin and microtubule concentration per cell by increasing seeding density. The increase in microtubule mRNA was higher than F-actin, whereas in the polymerized state (image-based concentration measures) the increase in F-actin was higher than microtubule.

### Summary and Correlation Analysis

In summary, model embryonic MSCs adapt their size, shape, cytoskeleton and stiffness in response to both mechanical cues comprising local compression induced by seeding at increasing target densities as well as stabilization of microtubules via exogenous PAX exposure (Table 1). In context of this experimental model system where exposure to PAX is known to stabilize polymerized microtubules by decreasing depolymerization, the 0.411 linear correlation coefficient between per cell microtubule concentration and PAX concentration can be considered as strongly correlated and other correlation values can be considered relative to that known strongly positive correlation. Interestingly, MSCs’ functional mechanoadaptation via stiffness modulation shows a higher, significant correlation to increasing local compression (seeding density, correlation coefficient 0.415) than to exogenous microtubule stabilization (PAX, correlation coefficient 0.250). In contrast, MSC’s morphological adaptation via cell volume and shape, nucleus volume and shape and microtubule concentration show contrasting correlations to exogenous PAX exposure compared to local compression via increased seeding density. Whereas native (no exogenous PAX exposure) MSCs show highest correlations for adaptation via modulus stiffening, microtubule stabilized (PAX exposed), enlarged MSCs exhibit highest and similar correlations to cytoskeletal concentrations, e.g. of microtubules and actin.

**Table 1.** Correlation matrix relating independent variables, PAX exposure and local compression to cell mechanoadaptation variables (first two rows). The correlation coefficient is shown for pooled data and indicates the level of interaction between cytoskeletal adaptation/remodeling parameters. Green fields indicate a positive correlation, pink fields indicate a negative correlation, and gray fields indicate no correlation, with cell values indicating Pearson’s correlation coefficient. *Correlation significant at the 0.05 level (2-tailed). **Correlation significant at the 0.01 level (2-tailed).

| Correlations |  |  |  |  |  |  |  |  |  |  |
| --- | --- | --- | --- | --- | --- | --- | --- | --- | --- | --- |
|  | PAX concentration | Seeding density | Cell volume | Cell shape | Nucleus volume | Nucleus shape | Actin concentration | Microtubule concentration | Actin alignment | Youngs modulus |
| PAX concentration |  | -0.005 | 0.501** | 0.403** | 0.575** | 0.635** | 0.415** | 0.411** | 0.26 | 0.250** |
| Seeding density | -0.005 |  | 0.163** | 0.233** | -0.147** | 0.046 | -0.320** | -0.218** | -0.408* | 0.415** |
| Cell volume | 0.501** | 0.163** |  | 0.066 | 0.212** | 0.398** | 0.211** | 0.418** | -0.203 | 0.298** |
| Cell shape | 0.403** | 0.233** | 0.066 |  | 0.186** | 0.355** | 0.131* | 0.134* | 0.219 | -0.123 |
| Nucleus volume | 0.575** | -0.147** | 0.212** | 0.186** |  | 0.316** | 0.297** | 0.198** | 0.134 | -0.016 |
| Nucleus shape | 0.635** | 0.046 | 0.398** | 0.355** | 0.316** |  | 0.284** | 0.332** | 0.142 | -0.009 |
| Actin concentration | 0.415** | -0.320** | 0.211** | 0.131* | 0.297** | 0.284** |  | 0.336** | 0.399* | -0.093 |
| Microtubule concentration | 0.411** | -0.218** | 0.418** | 0.134* | 0.198** | 0.332** | 0.336** |  | 0.458** | -0.05 |
| Actin alignment | 0.26 | -0.408* | -0.203 | 0.219 | 0.134 | 0.142 | 0.399* | 0.458** |  | -0.454* |
| Youngs modulus | 0.250** | 0.415** | 0.298** | -0.123 | -0.016 | -0.009 | -0.093 | -0.05 | -0.454* |  |

## Discussion

Below a threshold concentration of 100nM, exposure of model murine embryonic stem cells to the microtubule stabilizing agent PAX diminishes depolymerization of the tubulin cytoskeleton and perturbs the intrinsic mechanoadaptation capacity of the cells while maintaining cell viability. Exposure to PAX stabilizes not only microtubules but also modulates actin de-/polymerization dynamics and spatial distribution (anisotropy and emergent architectures), as measured by increased F-actin alignment concomitant to significant increases in stiffness (Young’s Modulus) of in both adherent and suspended cells. In combination with increased local compression induced by seeding at increasing densities, PAX-exposed cells increase in cell volume with increasing PAX concentration, and with increasing significance at later time points. Hence, up to a threshold concentration of 100 nM, and in combination with seeding at higher density protocols to induce local compression, PAX provides an exogenous means to probe MSC mechanoadaptation under controlled mechanical loading conditions.

The use of molecular agents that target the de-/polymerization of F-actin and microtubule to modulate stem cell structure and function has been reported over the past several decades^7,10,20^. This has contributed to understanding the cytoskeleton’s regulatory role during lineage commitment^7,21–23^. PAX stabilizes the microtubule by binding to the β-tubulin subunit while at the same time allowing microtubule polymerization even without the presence of exogenous GTP^24^. In the current study, this stabilization was grossly evident in spatiotemporal microscope images of tubulin cytoskeletal distribution.

As a common anticancer agent, PAX targets highly metabolically active and dividing cancer cells, where at specific concentrations (IC50 of 2 – 5 nM^25^) abnormal mitotic progression and cell cycle arrest occur^26,27^. Due to stem cells’ known relatively higher resistance to PAX^22,23^, μM concentrations can provide nuanced control of cytoskeletal structure while ensuring stem cell viability^23^. In human mesenchymal stem cell (hMSCs), at concentrations of 300 nM, PAX maintains cell viability but reduces proliferation and migratory speed^7,22^ as microtubules continue polymerizing^23^, thus generating a larger pushing force that influences the cells’ mechanical properties^28^. Hence, the current study’s definition of relevant PAX concentrations to modulate microtubule-controlled cell structure and function, e.g. explicating PAX concentration-dependent relationships for stem cell size, shape, stiffness, metabolic and cytoskeletal remodeling activity, provide novel reference data for mechanistic studies of stem cell mechanoadaptation. In this way, i*n vitro* exogenous exposure to defined concentration ranges of PAX provides a tool to help decipher the concentration-dependent mechanical regulation of cytoskeleton that correlates to cellular structure-function adaptation.

The continued presence of stabilized microtubules polymerizing to form bundles persistently influences cell tension. For maintenance of adhesion and sensing of substrate mechanical properties, cells regulate the establishment of focal adhesion (integrin) and generation of traction through modulation of F-actin polymerization and myosin II activity^29^. During development, F-actin and microtubules adapt to the ever-changing fluctuation of traction force over time. The de-/polymerization of the F-actin and microtubule regulates cell shape and movement and generates forces balancing tension at cell boundaries^30^. F-actins’ contractile forces are partly counterbalanced by microtubules’ compression-resisting forces and the traction forces exerted onto the ECM, resulting in a cellular force balance or tensional homeostasis^31^. Hence in the current study, due to the cooperative function of F-actin and microtubules, the stabilized microtubule bundles lead to the increase in F-actin polymerization in a linear fashion that contributed to the distinct cell orientation and increased cell stiffness.

However, as noted above, the mechanisms by which the cytoskeleton maintains cell structure and associated function, and the mechanical properties of the resultant tissues are poorly understood. This provides the impetus for the current study and a follow on [Part B] study to map the adaptive cell shape and the cytoskeletal remodeling in response to controlled changes in the cell’s local environment and/or in the de-/polymerization dynamics of F-actin or tubulin cytoskeleton (Supplementary video 1). Upon stabilization, microtubule polymerization continues to generate pushing forces towards the cell periphery, the accompanying changes in F-actin may explain how cells distribute forces and adapt their structure. PAX exposure results in stem cell volume increase in a concentration– and time-dependent manner that correlates with increase in F-actin concentration, linear unbranched F-actin growth, and a high degree of alignment at the length scale of the cell. Considering the crosstalk between F-actin and microtubules, the presence of linear unbranched, highly aligned F-actin exerts additional control over microtubule gliding, resulting in further slowing down of microtubule movement^32^. As reported in *in vitro* reconstituted filaments, unbranched F-actin coexists with stabilized microtubules which in this case could be a more energetically favorable mechanism for the cells rather than allowing otherwise the ATP-required microtubule gliding force and movement.

In addition to time– and concentration-dependent volume increases and cell flattening associated with exogenous microtubule stabilization, stem cell nuclei also exhibit a concentration-dependent volume increase that is independent of time. Together, this presents an opposing shape shift between the cell and nucleus, i.e. microtubule stabilized cells exhibit a maximum increase in volume thus maintaining a small SA/V (indicative of rounder cells) while their nuclei exhibit an increasing SA/V or flattening over time. Cells maintain volume through mechanical constraints that render additional tension through the cytoskeleton, but the nucleus volume is regulated mainly by the volume of the cytoplasm and not mechanical constraints^33^. In the current study, control cells were allowed to proliferate, and the increase in cell number presents the mechanical constraint at cell boundaries that limits the cells’ volumetric growth. With microtubule stabilization, cell proliferation is low, and the stiffness of glass substrates provides the cues for polymerizing microtubules, leading to a maximum increase in cell volume. Polymerizing microtubules provide a greater compressive load within the cell that needs to be balanced by tension via F-actin, hence resulting in increased F-actin polymerization and contributing to a larger cytoplasmic component that in turn influences the growth in nuclear volume^34^. Additionally, microtubule stabilization deforms the nuclear membrane, reducing nuclear lamina proteins^30,35^ and causing fragmentation of the nucleus which over time results in increased nucleus surface area. The opposing shape changes between cells and their nuclei suggest that nuclei are still able to control their volume proportional to the total cell volume.

Mechanical constraints from nuclear crowding and spatial confinement also limit cell proliferation in terminally developing tissue^36^. Similarly, when higher seeding density was introduced to emulate the local compression in condensing mesenchyme, the PAX concentration-dependent cell volume increase was suppressed. In contrast, the nucleus volume increase could still reach maximum at a similar rate across the seeding densities over time. As a result, cells seeded at higher density exhibit decreasing SA/V (rounder) but their nuclei show increasing SA/V (flatter) with increasing PAX concentration. The way density is achieved also greatly influences cell morphometrics, whereby cells proliferating to high density exhibit rounder nuclei that correlate with significant upregulation of mesenchymal condensation gene markers^13^. With increasing density, cells seeded at target density exhibit flatter nuclei and greater number of overlaps with neighboring cells that correlate with less than two-fold up-or down-regulation of mesenchymal condensation markers^4,13^.

A recently reported FRET tool for the measurement intercellular tension generated by F-actin revealed that tension accumulates in stress fibers that experience stretch in the direction parallel to the fibers, whereas relaxation of fibers occurred when they experienced stretch in the perpendicular direction to them^37^. Such mechanical anisotropy of F-actin provides insights on the tension that F-actin may generate to accommodate the PAX-stabilized microtubules that at the same time guide the directionality of the fibers, including the radially arranged microtubules, and the cells overall polarity. The anisotropy of F-actin and microtubules was also observed in the level of increase in concentration per cell where the increase in F-actin was more significant in the basal region while the increase in microtubule is more significant globally in both apical and basal regions. Interestingly, the increase in mRNA levels of tubulin was more significant than the increase in actin, even when experiencing higher compression from increasing seeding density.

All experimental studies exhibit intrinsic limitations, and understanding of these limitations is crucial for putting results in context of e.g. stem cells *in situ*, in living organisms. Firstly, with regard to the use of PAX as an exogenous chemical modulator of microtubule dynamics, within specific ranges of concentration probed in Part A of these studies, PAX provides an appropriate tool to probe the effect of specific, controlled perturbations to the cellular machinery on stem cell adaptation and nascent lineage commitment. In this context, the controlled impeding of microtubule depolymerization is more specific than e.g. the use of genetic knock downs, while also likely exerting effects on other cellular functions and processes. 100 nM PAX was chosen as the upper bound for concentrations reported in this study, as the compensatory and adaptation mechanisms necessary for survival of multicellular constructs were of particular interest in this study. Secondly, in cognizance of intrinsic limitations of the C3H10T1/2 murine model embryonic stem cell line, use of the cell line has advantages, e.g. does not show the phenotypic drift observed in the primary cells, as well as provides a generalizable model cell line without the inherent single individual sourcing limitations e.g. of human MSCs, and a library of published reference data^2,17,38,39^ using the same density and culture models which serve as a foundation for the current study. Thirdly, the culture platform in which cells are seeded at increasing density provides a fully three-dimensional environment for seeded cells designed to emulate that of cells *in situ* within developing tissue templates.

Furthermore, Increasing cell seeding density has been shown to impede cell cycle progression^40^, arresting cell division at smallest cell volumes^41^. Interestingly, high seeding density has been reported to rescue cell cycle progression associated with cells cultured on soft substrates (500 Pa)^40^. This interplay of substrate stiffnesses and cell seeding density on cell cycle progression, and its relation to cellular traction goes beyond the scope of the current and follow on [Part B] study and deserves further investigation. Finally, microtubule stabilized cells have been shown to adapt and enter cell cycle arrest to maintain function, possibly via a prolonged G2 phase^7^. Here we consistently observe sensitivity to PAX at 10-100 nM in C3H/10T1/2 cells. Although we expect that cells were able to progress into early mitotic phase and synthesize new microtubules, given microtubules’ larger structure compared to F-actin, it is expected that stabilization of microtubules leads to a reduced capacity to facilitate the pulling forces required for cytokinesis, thus leading to arrest and reduced proliferation^27^.

In summary, our study reports the modulation of cytoskeleton using a defined concentration range of PAX that challenges cellular proliferation and metabolism, while maintaining viability and allowing reorganization of F-actin and microtubules. We demonstrate that by emulating stresses in development, e.g. the local compression via increasing seeding density, one can probe the capacity of cells to achieve force balance via opposing shape changes and cytoskeleton reorganization. We also revealed the extent to which PAX affects cell volume, and F-actin and microtubule concentration increases, modulated by increasing seeding density. The consistent effect of local compression in reducing the PAX-induced increase in both microtubule and actin at mRNA and filamentous level, suggests that mechanical cues are ubiquitously transduced to provide feedback at the gene level, manifesting in structure and function, i.e. the amount of tension cells need to generate or compression they need to resist.

## Materials and Methods

### Cell culture and exogenous microtubule stabilization

The C3H/10T1/2 murine embryonic stem cell line (CCL-226) was used as a model for primary embryonic mesenchymal stem cells and cultured in basal eagle medium (BME) with 1% penicillin-streptomycin, 1% L-glutamate, and 10% fetal bovine serum (FBS), per previously published protocols^2,13^. Cells were expanded and passaged to less than P15. For imaging, cells were seeded in glass-bottomed 24 well plates or 35 mm dishes. For inducing local compression through increasing seeding density^4,13^, cells were seeded at 5000 cells/cm^2^ for low density (LD), 15,000 cells/cm^2^ for high density (HD) and 45,000 cells/cm^2^ for very high density (VHD).

Exogenous microtubule stabilization via PAX exposure was carried out on the day following seeding. PAX solution was prepared from the stock of 5 mg/ml in Dimethyl Sulfoxide (DMSO) (5855 µM) in culture medium at a concentration range of concentration (1 – 100 nM). Medium was removed from the well plates and PAX solution was added to the wells. Control cells were treated with medium containing only DMSO. Cells were incubated until used for assays or imaging at determined time points (24, 48, and 72h).

### Cell proliferation and metabolic activity (MTT) assay

C3H/10T1/2 were seeded in 96-well plates at a density of 2000 cells per well (n=6). To determine the concentration-dependent response of MSCs to cytoskeleton targeting agents, on the following day, cells were exposed to paclitaxel (PAX) with a concentration range of 1 – 100 nM or Cytochalasin D (Cyto D) at 0.5 – 2 μM. At 16, 24, 48, and 72 h post PAX or Cyto D treatment, medium from plates was removed and cells were washed once with PBS. For CyQuant proliferation assay (Invitrogen), 100 µl Cyquant solution (22 µl Cyquant reagent A in 11 mL 1x Hanks’ Balance salt solution (HBSS) was added to each well. The plates were incubated in the dark for 30 minutes and then read for fluorescence using a Victor 3 microplate reader (Perkin Elmer) with excitation/emission detection at 485 nm/530 nm. For the MTT metabolic assay, MTT (3-[4,5-dimethylthiazol-2-yl]-2,5-diphenyltetrazolium bromide) solution was prepared by diluting the stock solution (5 mg/ml MTT in PBS) in culture medium at 1:10 dilution. 200 µl of MTT solution was added to each well and plates were incubated in 37C for 3 hours. The MTT solution was then removed, and the purple formazan was solubilized in DMSO. The absorbance was read using the plate reader at 570 nm.

### Labelling of cells, nuclei, and the cytoskeleton for confocal imaging

For analysis of live and dead cells, as well as measurement of cell and nuclei shape and volume, at pre-determined time points, cells were labelled with calcein AM (2nM) to indicate live cells and ethidium homodimer (EtHD-1) (4nM) to indicate dead cells. The nuclei were labelled with Hoechst (1μl/ml medium). After medium was removed, staining solution was added to the wells and cells were incubated for 30-45 minutes. Cells were then retrieved, washed with PBS twice and covered with fresh medium. The cells in well plates were then taken for confocal imaging.

For analysis of cytoskeleton structure, cells were fixed at pre-determined time points after PAX exposure with 4% paraformaldehyde for 10 minutes, permeabilized with 0.1% Triton™ X-100 in 1X PBS, and blocked with 10% FBS in PBS, with 3 times PBS wash in between. Cells were blocked for unspecific binding using 5% bovine serum albumin (BSA) in PBS for 1 hour. Then, tubulin was labelled with α-tubulin monoclonal antibody 2 µg/mL (Life Tech) for 3 hours at room temperature then crosslinked with secondary Goat anti-Mouse IgG conjugated with Alexa Fluor 568. For labelling F-actin, cells were stained with ActinGreenTM (AlexaFluor™ 488 phalloidin).

### Confocal imaging and analysis of cell and nuclei shape and volume

Imaging was performed on Leica SP8 confocal microscope. Prior to imaging, cells were stained with Hoechst for nuclei, 2 µM calcein AM for live cells, and 4µM ethidium homodimer (EtHD-1) for dead cells and incubated in the dark for 1 hour. Nuclei were imaged using diode 405 nm laser line. EthD-1 labelled cells were imaged using 561 nm helium neon laser line. Calcein AM labelled live cells were imaged using the 488 nm argon laser line. A sequential scan was performed on all fluorophores between frames, with each scan taken at 400 Hz. A 40x, 1.3 NA oil immersion objective lens was used for imaging. The imaging format was set at 2048 x 2048, resulting in an image size of 387.69 μm^2^ and pixel size of 0.189 μm. 3D images were obtained by scanning in z direction the whole cell thickness (average of 10 μm thick) with 0.5 µm step size, giving the average of 18-20 slices per 3D image.

Geometry of cells and their nuclei was quantified, including measures of volume, surface area, and the surface area to volume ratio (SA/V). These analyses were done using a custom MATLAB code that allows semi-automated analysis on selected single cells in each Field of View (validated in comparison with IMARIS, Fig. S5). The code first converted the 3D images into binary images, then applied masks on the selected individual cells from each image. A threshold was then applied to exclude noise. Statistics models were applied to parameters including volume, surface area, and principal axes lengths (x, y, z), and convex volume. The corresponding data describing each voxel, of whole cells and nuclei, were then extracted for statistical analysis.

## Statistical analysis

Statistical analysis of experimental data was performed using Graphpad prism (La Jolla, CA) and SPSS (IBM). Significant differences in cell volume, stiffness, F-actin and microtubule concentration across PAX concentration, substrate stiffness and seeding densities were analyzed with Two-way ANOVA and Tukey’s multiple comparison test. Linear regression was performed to confirm concentration-dependent effect of PAX on cell volume (Fig. 6). Interactions between the independent (PAX concentration and seeding density) and outcome variables (cell volume, cell shape, nucleus volume, nucleus shape, actin and microtubule concentration, actin alignment and Young’s Modulus) were tested using a Correlation Analysis (Table 1).

### Immunofluorescence staining and confocal imaging of cytoskeleton alignment

Imaging of cytoskeleton labelled cells was performed on Zeiss 900 confocal microscope with Airy scan mode. Hoechst labelled nuclei, phalloidin labelled F-actin and immunofluorescence labelled α-tubulin were imaged using the respective 405 nm, 488 nm and 561 nm diode lasers. Super resolution (SR) acquisition mode was selected which automatically gave a pixel size of 0.05 µm. Zoom factor was set at 0.5x to give an image size of 202.8 µm x 202.8 µm and frame size of 3592 px x 3592 px. A 63x, 1.4 NA oil immersion objective lens was used for imaging. Scan speed was set at 35.55s. 3D images were acquired by selecting z-stack mode with step size of 0.25 µm.

3D fluorescent images of F-actin were analyzed using a custom Local Orientation Image J plugin which operates and detects filaments using the Noise-based segmentation (NoBS) method, previously developed for neuronal alignment and described in further detail in the Supplementary Appendix (SA)^42^.

### Atomic Force Microscopy (AFM)

Cells were seeded in 35 mm glass bottomed dish (Fluorodish, WPI) at density of 5000 cells/cm^2^ and exposed to exogenous microtubule stabilization (PAX) for 24-72 h. Cells were washed with PBS and culture medium was replaced with serum free and phenol red free medium prior to AFM measurement. Both imaging and indentation were done using The JPK BioAFM (Bruker) mounted on Nikon inverted microscope connected to stage heater set at 37°C within a TMC vibration isolation table (Technical Manufacturing Corporation, MA, USA). Calibration of AFM system was done each time before the measurement to check the deflection sensitivity of the probe in liquid by engaging the probe on to the free glass surface, retraction of the probe and thermal noise calibration is to determine the spring constant.

A V-shaped cantilever with a pyramidal Scan Assist fluid tip probe with a nominal tip radius of 15 nm, a spring constant (k) of 0.7 N/m and a resonance frequency of 120–180k Hz was selected and mounted on the cantilever holder of the AFM scanner (Bruker)^43^. The Elastic modulus was determined by indentation in Contact Force Spectroscopy mode with pre-determined loading force and scan rate. Live cells were maintained in serum free and phenol red free medium during the measurement which was conducted no more than 2 hours to preserve cell viability. Cells with similar morphology that were not overlapping with other cells were selected for measurement and 10-15 force curves were acquired per cell on different location of the cell. The Young’s modulus values were extracted from force curves using Hertz and Sneddon fit and JPK analysis software.

### Real-time deformability cytometry

Cells were seeded in T-25 flasks at a density of 5000 cells/cm^2^. Cells were exposed the next day to either 100 nM PAX or DMSO (control). After 48h, the cells were harvested and centrifuged. The cell pellet was washed with PBS and resuspended in 1 mL CellCarrier buffer, PBS containing less than 1% methyl cellulose for adjusted viscosity. The cell suspension was then loaded into the syringe and flushed through the 30-μm narrow channel constriction in a microfluidic chip (AcCellerator, Zellmechanik Dresden). A total flow rate of 0.160 μl/s (0.04 μl/s sample flow and 0.12 μl/s sheath flow) was applied. Individual cells passing the channel were captured by high-speed camera from which cell contour were tracked and information such as cell area (A), deformation (D) could be derived using the formula D=1-2√πAl, where l = perimeter of the contour. A bright field image was acquired for every measured cell, allowing multiparametric offline analysis to discriminate single cells characteristics in a population. Data analysis and computation of the young’s modulus was performed in Shape-Out 2 software (available at https://www.zellmechanik.com/Download.html).

## Acknowledgments

The authors would like to acknowledge the Katharina Gaus Light Microscopy Facility (KGLMF) and their staff for the continuous support in imaging resources and assistance in image processing and analysis. Special acknowledgement to Dr. Michael Carnell for developing the NoBS algorithm for the analysis of F-actin stress fibers alignment; Dr. Elvis Pandzic for assisting and training V.P. in custom-built MATLAB scripts for the analysis of cell shape, volume; and Dr. Celine Heu for assistance with the AFM; Dr. Chantal Kopecky for assistance in deformability cytometry. V.P. would like to thank the Scientia PhD Scholarship scheme from UNSW for support throughout her doctoral research studies. This work has been supported in part by grants from the National Health and Medical Research Council (MLKT, KAK) and the National Institute of Health (KAK).

