## Supplemental information and figures for "Stem cell mechanoadaptation - Part A - Effect of microtubule stabilization and volume changing stresses on cytoskeletal remodeling"

**This PDF file includes:**

Supporting text

Figures S1 to S5

SI References

**Supporting Text**

**Detailed Noise Based Method Segmentation (NoBS) Method**

First, NoBS algorithm differentiates objects and background noise through overestimation of objects that include *circa* 50% of background pixels, to effectively detect faint details of filaments. The overestimation uses 24 binary images first created from the original intensity image containing object filaments emanating in various orientations and distance within the local 7x7 pixels. A positive label is given to a pixel if its value is greater than the average of its relevant flanking neighbors (pixel pairs) for its specific angle and distance. Later, false positive background pixels are suppressed by applying filters (Filter 1) comprising a human-trained pattern detector that observes the local 5x5 pixels and compares them to the previously created 24 image patterns that distinguish objects and noise. The 24 denoised binary images were combined as a single image and then used as a second human-trained filter (Filter 2) or reference pattern to further suppress noise. This method retains the weak signals that may have only been present in a few scales and/or angles and are adjacent to each other spatially, whilst also suppressing objects that appeared to be detected in random angles or distance in many of the 24 images. Suppressed, weak, and strong patterns are all processed through Filter 2 which then results in images of strong or weak structures only. In this study, all strong structures are preserved and only weak structures that are connected with the strong structures are kept, thereby improving filament connectivity.

Operation of filters: Filter 1 was created using a set of mixture of fluorescent labelled F-actin filaments images reserved for training purposes, as well as an additional computer-simulated data that was designed to resemble these images. Examining all 24 neighboring pixels in the 5x5 neighborhood in a binary image gives 224 (16777216) possible patterns, assuming the centre pixel is positive. The filter was initially set for all patterns to be suppressed. Using this set of training images (n > 20), patterns present in objects were gradually deselected. Training speed was enhanced by automatically including rotations and mirror-images of patterns which had already been deselected. 1,293,807 patterns were trained as objects, whilst 14,483,409 remained as noise. The same set of training images were used to create Filter 2 which again interrogates the 24 neighboring pixels and compares for their angles and distance, then gives the possible 224 patterns. Simulated data included images of lines, circles, and puncta of varying widths with differing degrees of Gaussian noise added. A training program was created allowing users to hover over a pixel of interest and visualize angles and distances that are positive. The user can then include or suppress pixels based upon their specific patterns. Training speed was again enhanced by including rotations and mirror patterns of those which were selected.

**Additional Discussion Points related to Seeding Cells at Density to Achieve Local Compression**

In previous studies, the density and developmental context of seeded stem cells significantly affected cell volume but not cell shape^1,2^: “cell volume was found to decrease as cell density increased and was also found to decrease with time spent in culture to achieve the target density”. Interestingly, “cell volume did not differ between cells that were proliferated to achieve target density, indicating that over longer periods of time, initial cell density rather than actual cell density modulates cell volume^”1,2.^

Furthermore, cell shape, when normalized to cell volume, was shown not to change significantly in response to cell density or seeding conditions. The cell shape and volume data from the aforementioned previous study provided experimental evidence that "biophysical cues, induced through cell seeding density and protocol, modulate the volume of mesenchymal stem cells as well as the shape of their nuclei. Furthermore, based on cross correlation analysis with gene expression data from a previous study using identical cell seeding protocols, these changes in cell volume and nucleus shape correlate significantly with changes in expression of genes marking mesenchymal condensation and the commitment of mesenchymal stem cell fate, including osteogenic, chondrogenic and adipogenic differentiation. Specifically, seeding mesenchymal stem cells at increasing target density is associated with a concomitant decrease in cell volume, reaching a steady state, with time...”^3,4.^

**Figures S1 to S5**

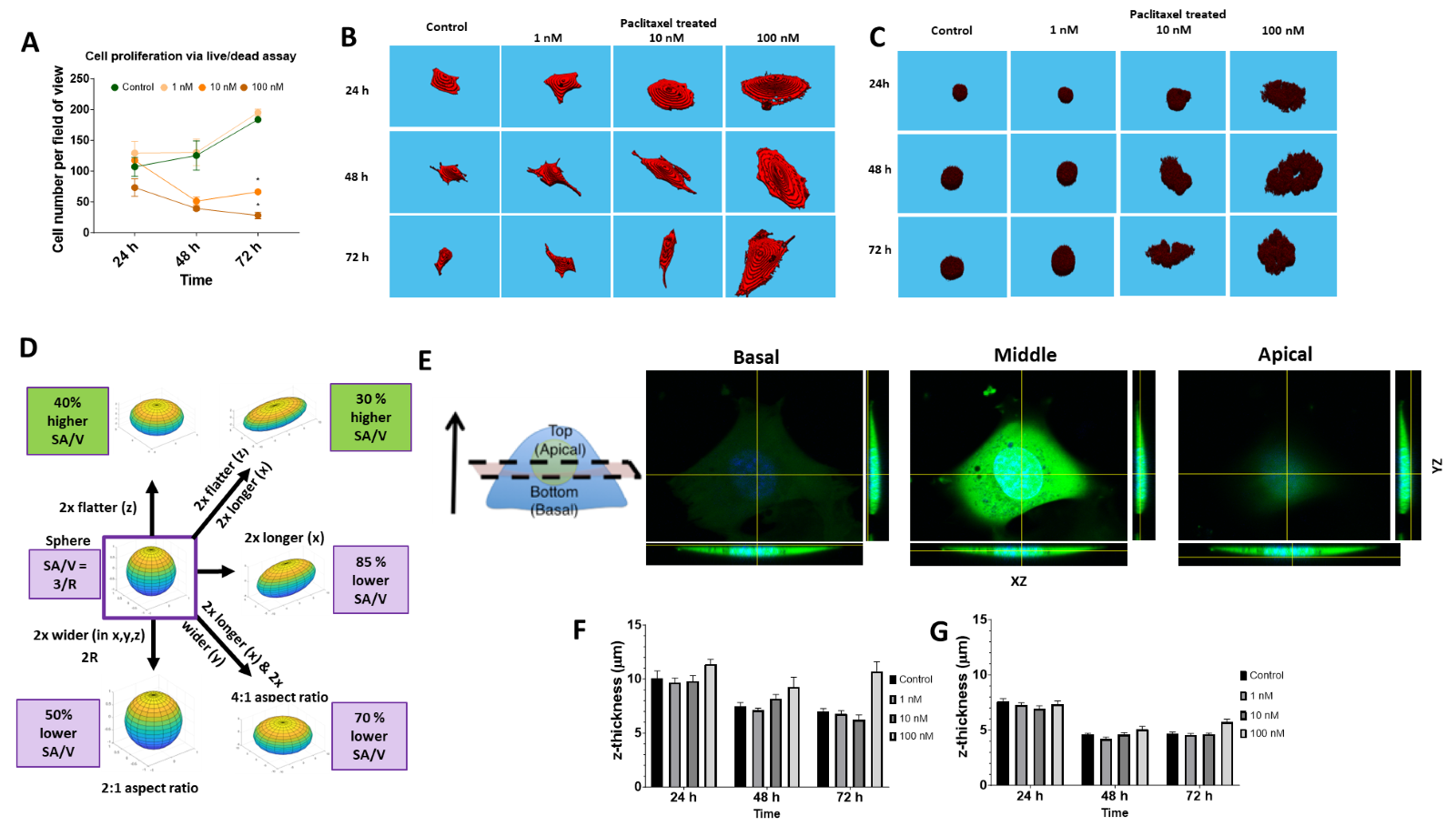

**Figure S1.** **Modulation of stem cell’s structure and function using microtubule stabilizing agent, Paclitaxel (PAX). (**A) Cell proliferation quantified by measure of cell number per field of view during imaging with live/dead assay, also shows structural adaptation as shown in the changes in cell and nuclei volume and shape. The masked binary single cell images from the custom-built MATLAB code are used to quantify the cell (B) and nucleus (C) volume upon treatment with PAX over time. (D) Cell and nucleus shape are defined as surface area to volume ratio (SA/V) and are normalized to the SA/V of a perfect sphere (3/R, R the cell radius) of the same volume. When cells or nuclei become rounder, their SA/V is close to 1 and when the cells or nuclei f latten their SA/V is greater than 1. (E) Schematic and the orthogonal view of cell and the division between apical and basal regions from the mid-plane of the nucleus. Calcein green represents cell and Hoechst (blue) represents nucleus. In general cell thickness (F) reduces over 72h in culture however cell thickness remains high at 100 nM PAX; in contrast, nuclear thickness (G) remains consistent in control cells and with PAX treatment.

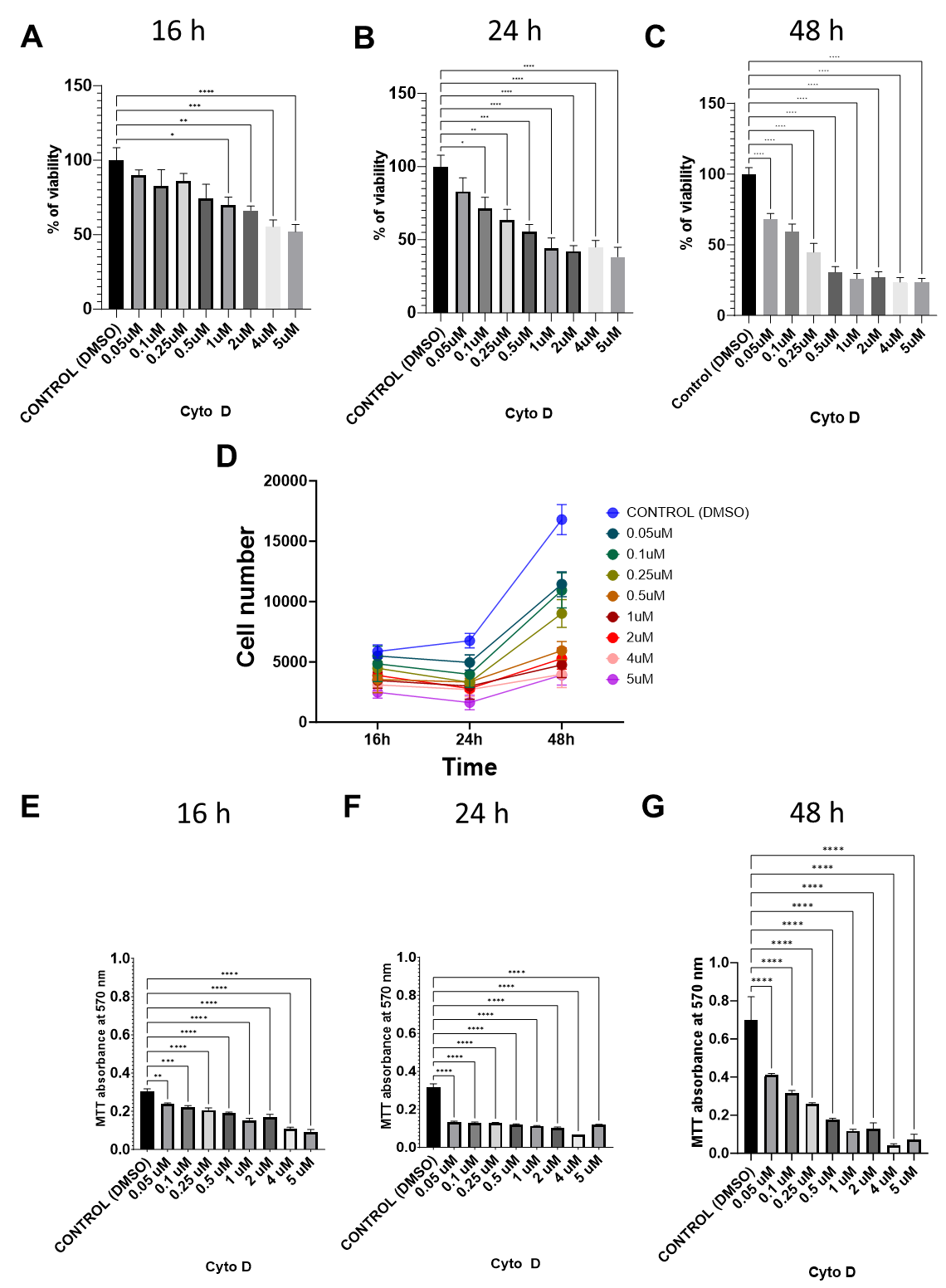

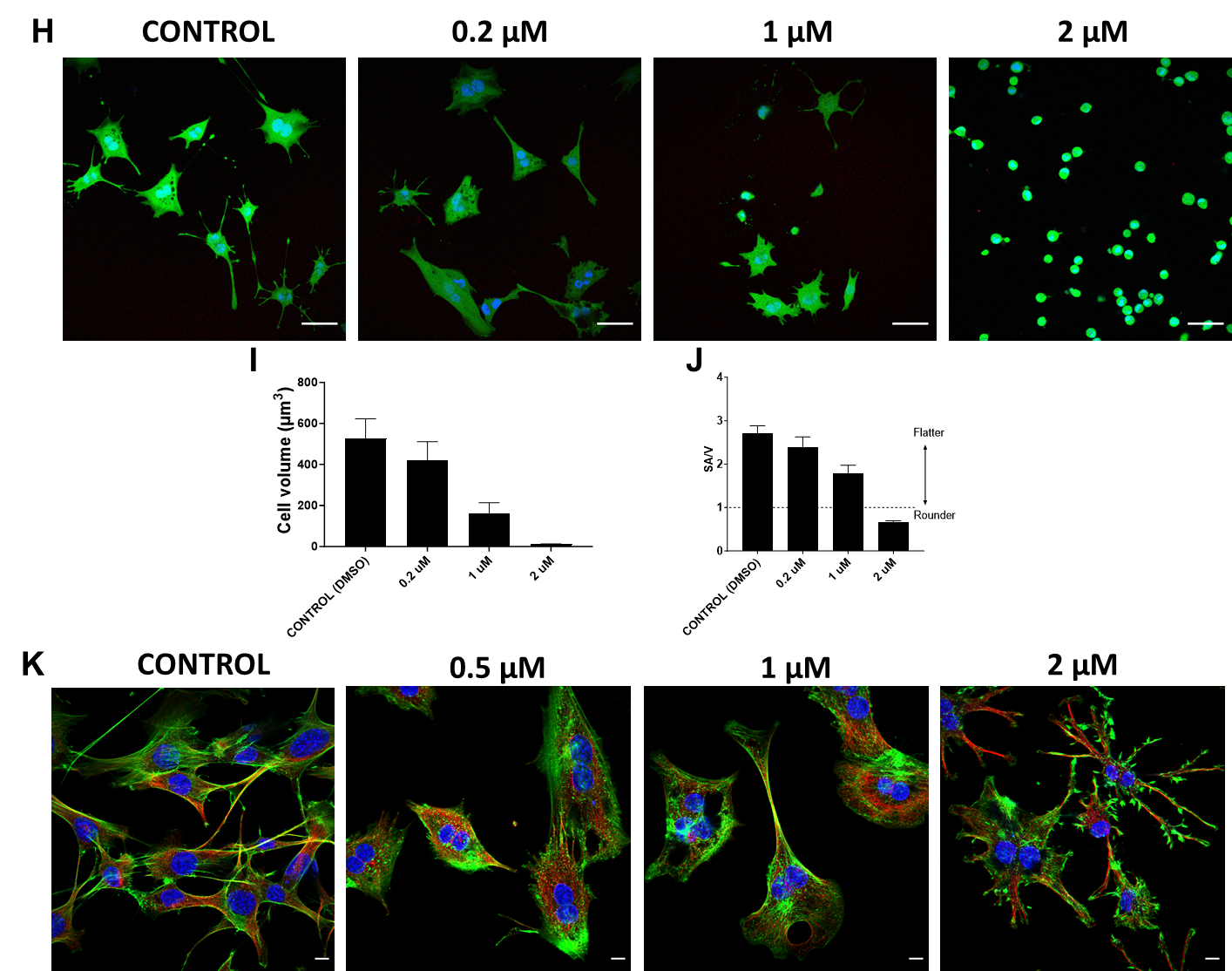

**Figure S2.** **Modulation of stem cell’s structure and function using actin stabilizing agent,** Cytochalasin D (Cyto D). C3H/10T1/2 viability at 16h (A) 24h (B), and 48h (C) measured with CyQuant assay. Standard curve of a known cell number from Cyquant readings were used to plot cell proliferation (D) over 48 h of Cyto D treatment. Cell number is calculated per well of 96-well plate. Metabolic activity measured via MTT assay af ter Cyto D treatment for 16 (E), 24 (F), and 48 h (G). (H) Live and dead assay (scale bar = 50 μm) reveal the immediate volume (I) and cell shape (J) changes with increasing cyto D concentration. Surface area to volume ratio (SA/V) was normalized to SA/V of sphere of the same volume. (K) Cyto D inhibits actin polymerization and removes the tension in the actin (green), resulting in thinner microtubule (red) bundles and shrinking cell morphology (scale bar = 10 μm).

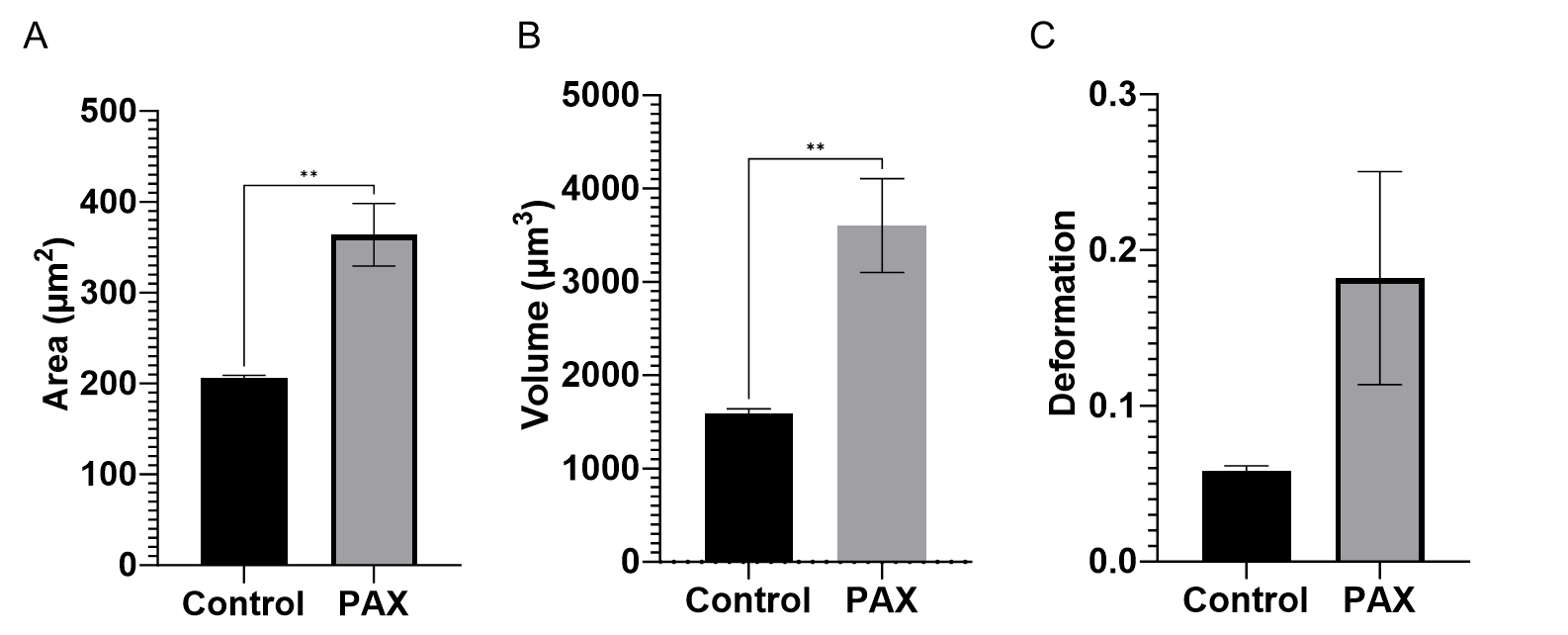

**Figure S3. The changes in the structure of PAX-treated MSCs harvested in suspension and flushed through the 30 μm width channel of the deformability cytometry.**

Treatment with PAX for 48h consistently increases the area (A) and volume (B) and deformation (C) of cells. (D) Treatment of MSCs with PAX results in increased cell volume. Top: Linear regression analysis of cell volume increase with increasing PAX concentration over 72h – showing that cell volume increases with increasing PAX concentration, and with increasing significance at later time points. Bands represent 95% confidence intervals. Error bars represent standard error of men (SEM). Bottom: Two-way ANOVA, fixed effect type III analysis of cell volume increase with increasing PAX concentration (based on Figure 2D in manuscript), with Tukey’s multiple comparison test, demonstrates that PAX-mediated cell volume increases correlate with time.

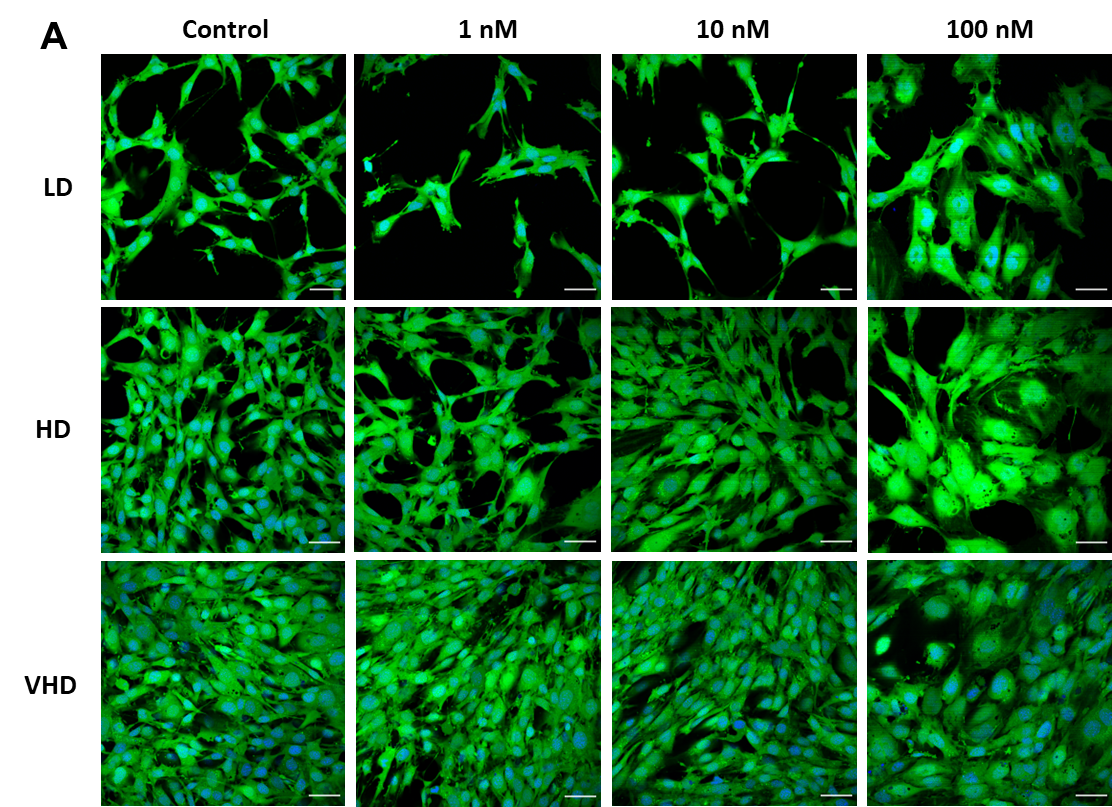

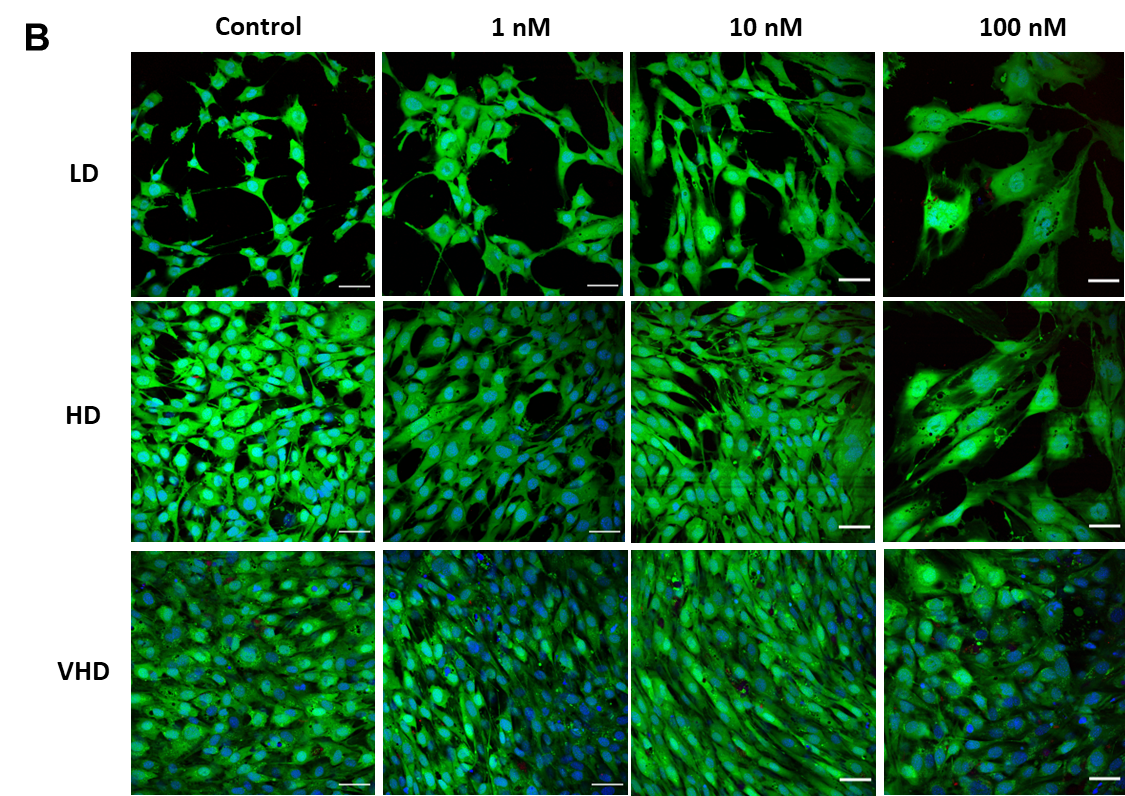

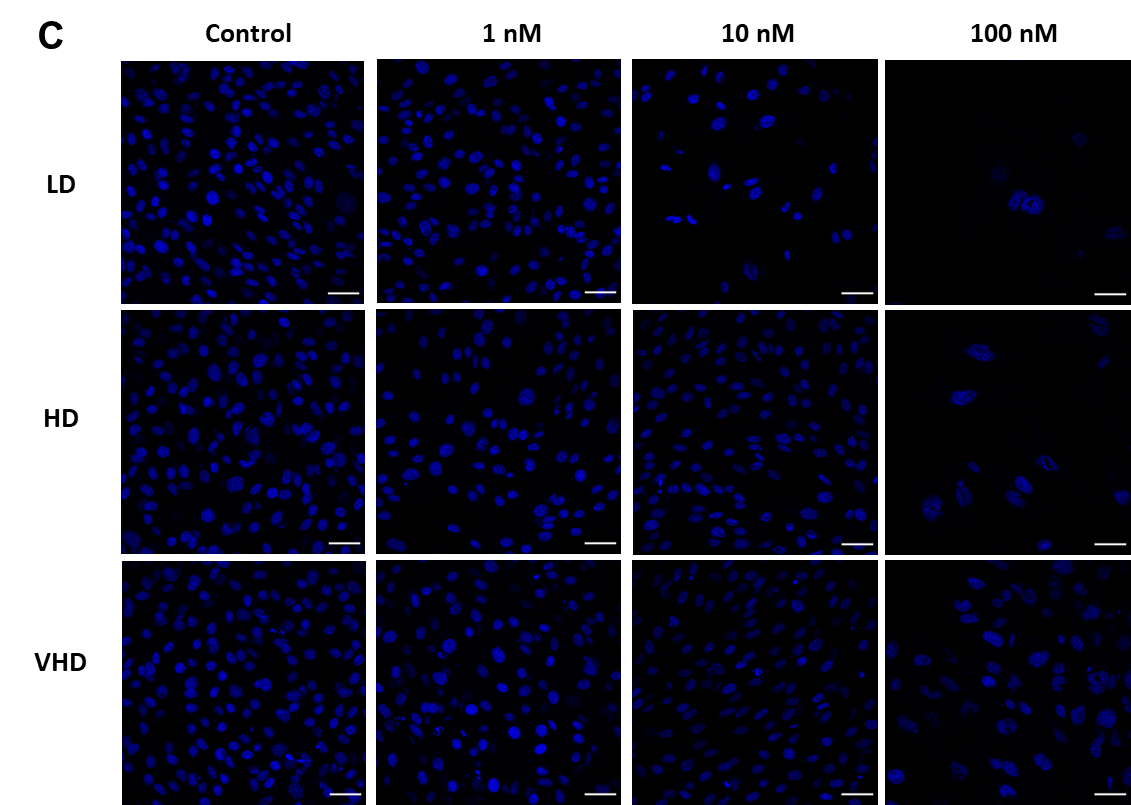

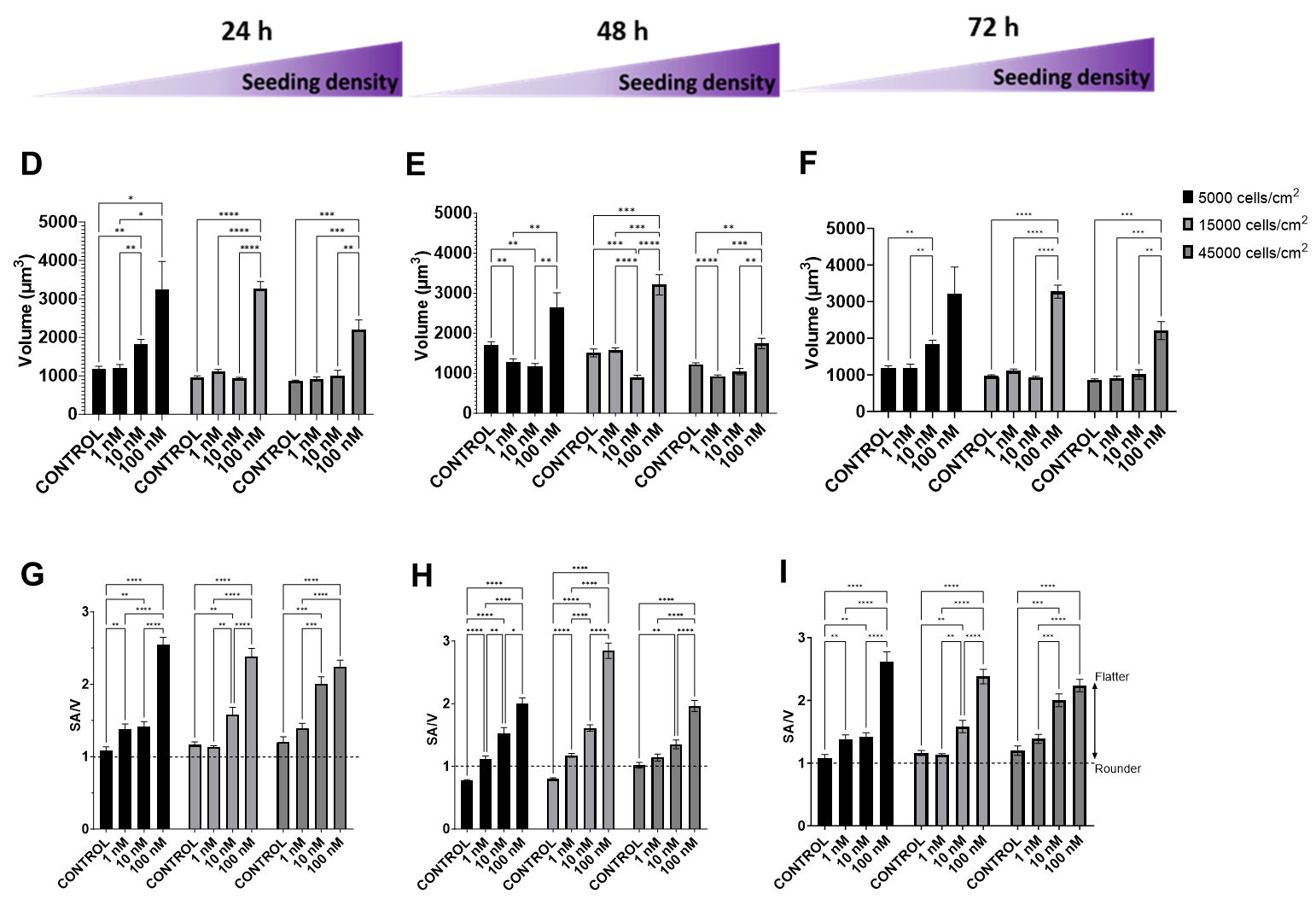

**Figure S4. The changes in volume and shape or surface to volume ratio (SA/V) of the nuclei upon culture of MSCs in various seeding densities as a means to introduce local compression.** The changes in cell shape and volume as a response to PAX and increasing seeding density at (A) 24 h and (B) 48 h of culture. (C) The nuclei exhibit morphological changes due to deformation by PAX. When seeded at density for 72h culture the nuclei volume seems to be dependent on PAX and cell seeding density, but not time. Across the 24h (D), 48h (E), and 72h (F) of PAX treatment, the nuclei volume increase due to PAX treatment is concomitant to cell volume increase (Scale bar = 50 μm). The increase in nuclei SA/V due to PAX is also comparable to the increase in cell SA/V across the seeding densities. The nuclei SA/V increase was similar in all time points 24h (G), 48h (H), and 72h (I), and not significantly modulated by increasing seeding density. Error bars represent ± standard error of mean. Significant differences are presented between neighbouring values from Tukey’s multiple correlation test (**** p < 0.0001, *** p<0.001, ** p < 0.01, * p< 0.05).

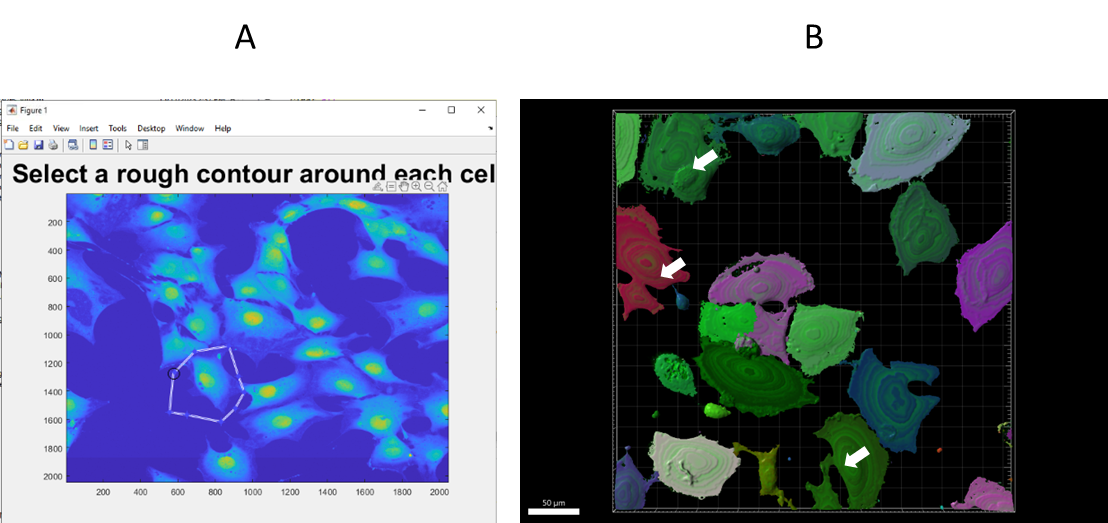

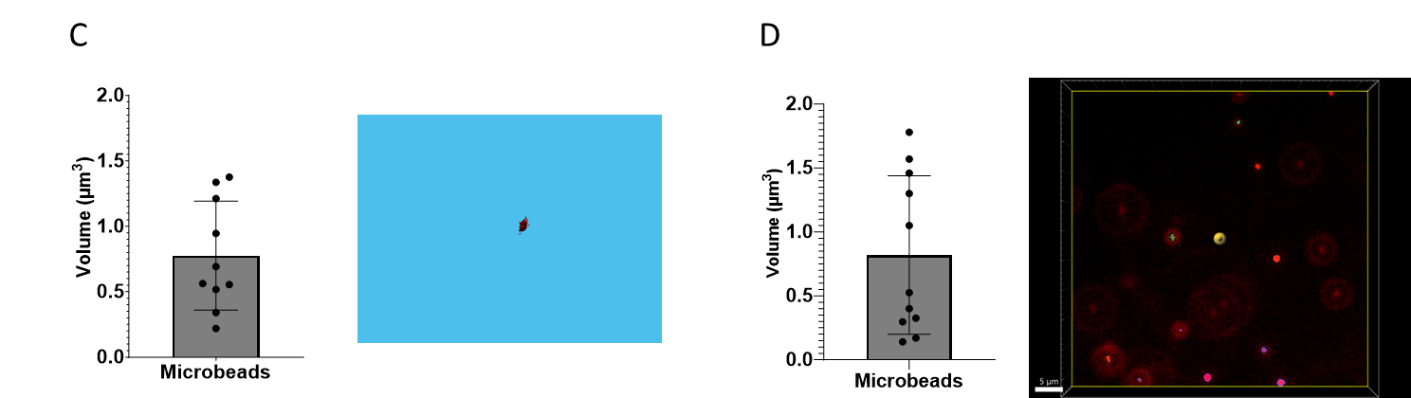

**Figure S5. Validation of cell volume quantification methods. (**A) The semi-automated image analysis using a custom-built MATLAB script enables the free drawing of cell contours to separate single cells. (B) Cell volume analysis using IMARIS is more automated, and the adjusted threshold is at times unable to separate clusters of closely adjacent cells (white arrows). (C, D) Volume analysis of commercially available polystyrene microbeads (BangsLab) ⌀ = 0.95 – 1.05 μm quantified using the custom-bulti MATLAB script (C) and IMARIS (D) validates the quantification methods in a controlled context.
